# Fasting reverses drug-resistance in hepatocellular carcinoma through p53-dependent metabolic synergism

**DOI:** 10.1101/2021.02.10.430545

**Authors:** Jelena Krstic, Isabel Reinisch, Katharina Schindlmaier, Markus Galhuber, Natascha Berger, Nadja Kupper, Elisabeth Moyschewitz, Martina Auer, Helene Michenthaler, Christoph Nössing, Maria R. Depaoli, Jeta Ramadani-Muja, Sarah Stryeck, Martin Pichler, Beate Rinner, Alexander J.A. Deutsch, Andreas Reinisch, Tobias Madl, Riccardo Zenezini Chiozzi, Albert J.R. Heck, Meritxell Huch, Roland Malli, Andreas Prokesch

## Abstract

Cancer cells voraciously consume nutrients to support their growth, exposing a metabolic vulnerability that can be therapeutically exploited. Here we show in hepatocellular carcinoma (HCC) cells, xenografts, and in patient-derived HCC organoids that fasting can synergistically sensitize resistant HCC to sorafenib. Mechanistically, sorafenib acts non-canonically as an inhibitor of mitochondrial respiration, causing resistant cells to depend on glycolysis for survival. Fasting, through reduction in glucose and impeded AKT/mTOR-signaling, prevents this Warburg shift. Regulating glucose transporter and pro-apoptotic protein expression, p53 is necessary and sufficient for the sorafenib-sensitizing effect of fasting. p53 is also crucial for fasting-mediated improvement of sorafenib efficacy in an orthotopic HCC mouse model. Together, our data suggest intermittent fasting and sorafenib as rational combination therapy for HCC with intact p53 signaling. As HCC therapy is currently severely limited by resistance, these results should instigate clinical studies aimed at improving therapy response in advanced-stage, and possibly even early-stage, HCC.

**HIGHLIGHTS:**

- Fasting sensitizes resistant HCC xenografts and patient-derived organoids to sorafenib
- Sorafenib-mediated Warburg shift is prevented by glucose limitation upon fasting
- Fasting synergistically improves sorafenib efficacy in non-resistant models
- p53 is required for synergism by regulating glucose uptake and apoptosis

**GRAPHICAL ABSTRACT:** 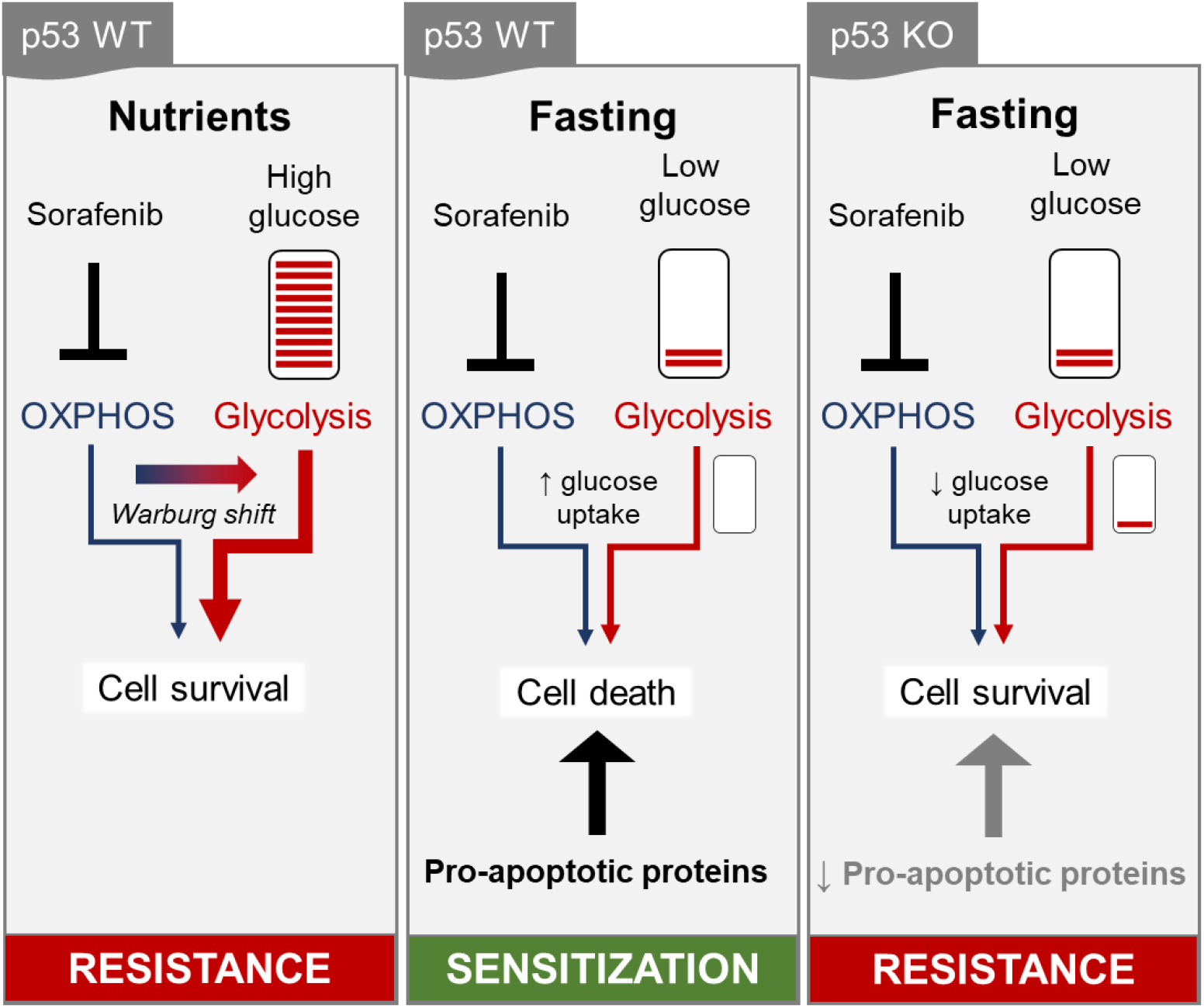

## Introduction

Many cases of advanced-stage solid tumors show only temporary responses to targeted therapy [1] due to the development of resistance against initially effective standard-of-care drugs. The ensuing residual disease [2] constitutes a severe impediment to progression-free patient survival. The adaptive nature [3] and metabolic flexibility of cancer cells [4,5] are key enabling mechanisms of resistance against targeted cancer therapy. Combination therapies represent promising strategies since they target several pathways, leaving less wiggle room for cancer cells to thrive on alternative growth-enhancing molecular routes.

Dietary interventions, in particular nutrient restriction regimens such as fasting or ketogenic diet, harbor a vast potential to support conventional cancer therapies [6–9]. The rationale underlying the use of fasting in cancer therapy is based on specific and general features of cancer cells: They disobey anti-growth signals [10] and possess a pronounced anabolic appetite [11,12]. Therefore cancer cells are unable to properly adapt to fasting conditions [13]. A large number of preclinical studies support this reasoning by showing reduced tumorigenesis, alleviation of therapy resistance, or reduced adverse effects when different fasting regimens are combined with standard-of-care drugs [13–18]. Several clinical trials are ongoing (reviewed in [19]) and will soon deliver data about the applicability and translatability of data from animal models. It is important to note, however, that the responses to fasting are specific to cancer types and their mutational landscapes [8]. Hence, focused studies need to validate fasting/drug combinations, factoring in cancer type and patient- as well as intra-tumor heterogeneity, to enable stratification approaches as recently suggested [6].

Hepatocellular carcinoma (HCC) is the primary form of liver cancer [20] and is one of the deadliest cancers worldwide with rising incidence [21]. This is due to the lack of response to classical chemotherapeutics (e.g. doxorubicin, cisplatin) and targeted drugs in early-stage disease [22,23]. For late-stage HCC, sorafenib has long been the mainstay in first-line treatment [24] and is still frequently used where other more expensive drugs are not available or upon progression after atezolizumab/bevacizumab in later treatment lines [25,26]. Sorafenib has been shown to act as multiple kinase inhibitor on the endothelial cell compartment (anti-angiogenic via e.g. VEGF inhibition) and on hepatocytes (mainly as RAF inhibitor) [27]. In a landmark placebo-controlled clinical trial sorafenib was shown to improve overall survival by three months with no case of complete remission [28]. This modest therapy success is largely ascribed to the development of sorafenib resistance [29,30]. While several mechanisms of resistance have been suggested, many reports converge on hyperactivation of PI3K/AKT/mTOR and MAPK signaling upon sorafenib treatment (reviewed in [31]). Thus, there is a need for novel treatment approaches and combination therapies that improve the current clinical situation for HCC patients.

Here, we show *in vivo* and *in vitro* evidence that nutrient restriction can sensitize sorafenib-resistant HCC models and improve the efficacy in sorafenib-responsive models. We pinpoint the mechanism of sensitization to a synergistic action of sorafenib, as a potent inhibitor of mitochondrial respiration, and starvation or fasting, which prevents a Warburg shift by limiting glucose availability. We further show that p53 is necessary for this sensitization by coordinating glucose uptake and pro-apoptotic protein expression. Taken together, our data suggest fasting as a potent adjuvant to sorafenib in p53-positive HCC and should inspire clinical studies that scrutinize this rational polytherapy approach.

## RESULTS

### Nutrient restriction synergistically sensitizes resistant HCC cells, xenografts, and patient-derived HCC organoids to sorafenib

Intrinsic and acquired sorafenib resistance is a major impediment to progression-free survival in late-stage HCC [27,31] and novel combination therapies to overcome resistance are needed. To investigate whether starvation can alleviate sorafenib resistance, we tested the responsiveness of HCC-derived HepG2 cells to sorafenib. Confirming previous studies [32], HepG2 cells stayed largely resistant to sorafenib up to supra-clinical doses (Figure S1A). In HCC patients, plasma sorafenib levels range between 5 and 20 µM after oral application [33,34]. Next, we subcutaneously injected these cells in NMRI nude mice and randomly divided them into four groups receiving either vehicle control or sorafenib and further in groups that were kept on *ad libitum* diet or on an intermittent fasting (IF) regimen (Figure 1A). For IF, mice were withheld food for 24 hours two times a week, with two- or three-days *ad libitum* refeeding in-between. While this protocol robustly elicited fasting-mediated changes in plasma metabolites (Figure S1B), mice were able to fully regain the weight loss during the refeeding days (Figure 1B and Figure S1C-D). Less gentle fasting protocols such as every other day fasting led to progressive weight loss (Figure S1E-F), rendering this regimen unsuitable for prolonged treatment protocols. Sorafenib or vehicle control was orally applied at the beginning of the refeeding phase at a concentration of 30 mg/kg, corresponding to weekly administered doses in patients [28] (toxicity also evaluated in pilot experiments, Figure S1G-H). Similar to our *in vitro* observation (Figure S1A), tumor growth was affected neither by sorafenib treatment nor by IF alone (Figure 1C-D). However, combined treatment over four weeks significantly stunted *in vivo* tumor growth (Figure 1C-D). Histological analyses showed no difference in the staining of the proliferation marker Ki67 (Figure 1E-F), while a trend to larger patches of necrotic cells was observed in the combination treatment group as compared to sorafenib alone (Figure 1G-H). These results indicate that IF and sorafenib synergistically alleviate resistance possibly through inducing cell death.

**Figure 1.**
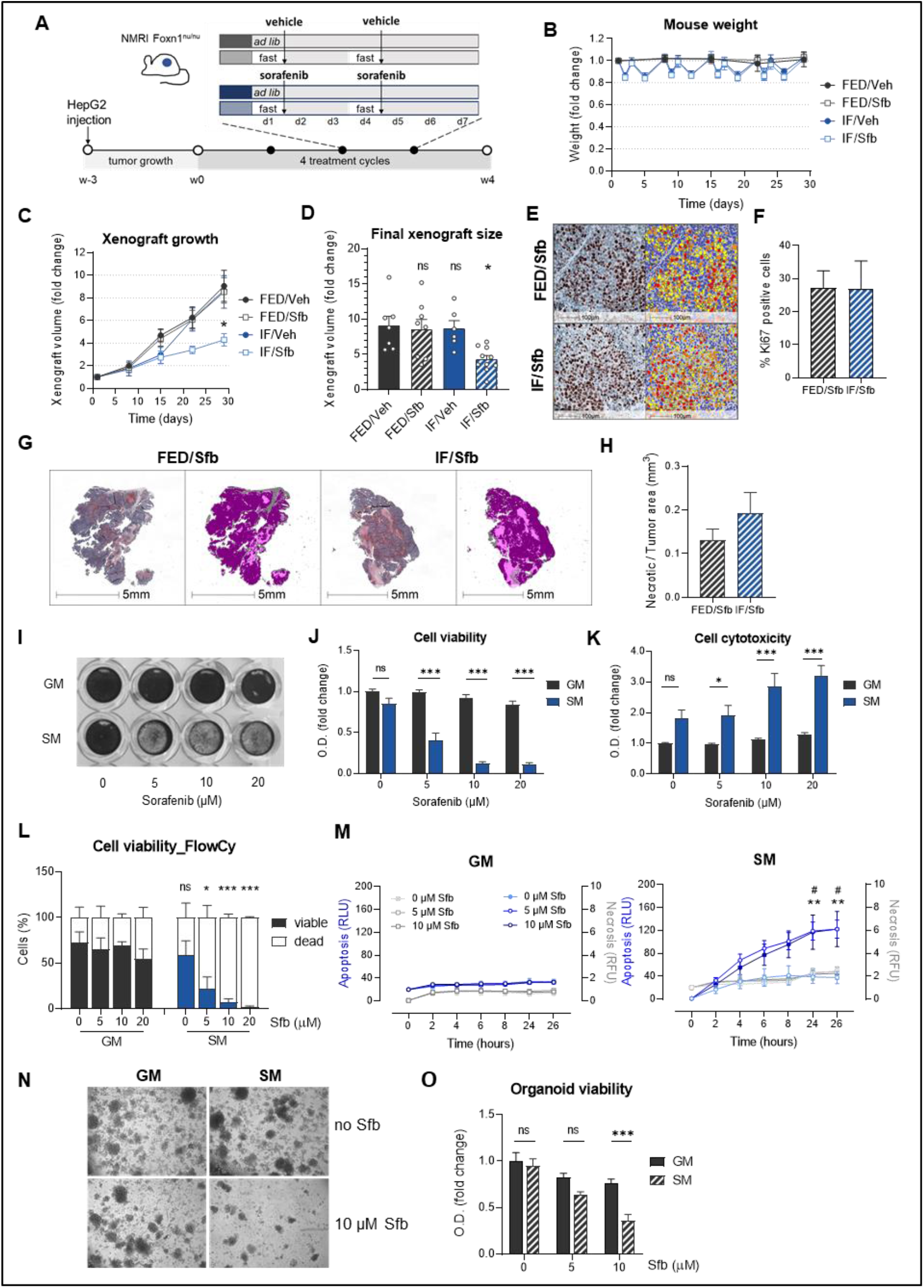
Nutrient restriction synergistically sensitizes resistant HCC cells, xenografts, and patient-derived HCC organoids to sorafenib. (A) Experimental design: HepG2 cells were subcutaneously injected into hind flanks of NMRI Foxn1nu/nu mice. After palpable xenografts were formed, mice were randomized into four groups: they were either fed ad libitum (FED) or intermittently fasted (IF) and received vehicle (Veh) or 30 mg/kg sorafenib (Sfb) twice per week for four weeks in total. (B) Animal weight during treatment, normalized to starting weight of each group. Mean values ± SD are shown; n=6 per group. (C) Xenograft volume normalized to the initial volume. All groups compared to FED/Veh group; two-way ANOVA, Dunnett’s multiple comparisons test, n=6-10 per group. (D) Relative xenograft volumes at the end of the experiment (day 29). Comparison of all groups vs FED/Veh group is shown; one-way ANOVA, Tukey’s multiple comparisons test; n=6-10 per group. (E) Representative histological images of xenografts from FED and IF mice treated with Sfb, labeled with antibody against proliferation marker Ki67. (F) Image analysis was performed by Halo® image analysis platform using Cytonuclear IHC module. Unpaired t-test was performed. (G) Necrotic area relative to tumor area quantified in histological xenograft sections (HE staining) from FED and IF mice treated with Sfb. (H) Image analysis was performed by Halo® image analysis platform using Random forest tissue classifier. Necrotic areas are colored pink. Unpaired student’s t-test was performed. (I) HepG2 cells stained with Gentian violet after 24 h of incubation in growth medium (GM) or starvation medium (SM) with 0, 5, 10 and 20 μM Sfb. (J) Viability of HepG2 cells after 24 h of incubation in GM or SM with indicated concentrations of Sfb. Values are normalized to the viability of non-treated cells grown in GM. (K) Cytotoxicity in cells grown in GM vs cells grown in SM with indicated Sfb concentrations after 24 h. (L) Percentage of viable (7AAD and Annexin V negative) and dead (7AAD and/or Annexin V positive) cells analyzed by flow cytometry. The comparison of SM groups to corresponding GM groups is shown. (M) Apoptosis (blue lines) and necrosis (grey lines) of HepG2 cells measured during 24 h of incubation in GM (left panel) and SM (right panel). Two-way ANOVA, Dunnett’s multiple comparisons test. The apoptotic/necrotic signal in each group was compared to control (GM/0). * SM/5 µM Sfb vs GM/0, # SM/10 µM Sfb vs GM/0. (N) (O) HCC patient-derived organoids were incubated in GM (expansion medium for HCC-organoids, see methods for formulation) or SM (consisting of 1/3 GM and 2/3 DMEM-/-/-, with final 5.5 mM glucose concentration) with or without Sfb for 6 days. (N) Microscope images showing a decreased number of organoids in combination treatment. (O) Organoid viability determined by viability assay. If not noted otherwise, mean values ± SEM are shown and two-way ANOVA, Tukey’s multiple comparisons test was performed. *** p<0.001, ** p<0.01, * p<0.05, ns-not significant (p>0.05); O.D. optical density.

To mechanistically investigate this synergism, we sought to mimic the nutrient restriction *in vitro*. For this, we used an HBSS-based starvation medium shown to induce autophagy in cultured liver cells [35] and to activate the fasting-responsive AMPK pathway in HepG2 cells [36]. When resistant HepG2 cells (Figure S1A) were kept in starvation medium for 24 hours, we observed a sorafenib dose-dependent reduction in viability, while starvation alone did not reduce viability (Figure 1I-J). These effects were exaggerated when viability was determined after 48 hours of treatment (Figure S1I-J). Furthermore, cytotoxicity (Figure 1K) and apoptosis (Figure 1L and S1K) were increased in a sorafenib dose-responsive manner in starved cells but not in cells kept in full growth medium. To gain insight into the kinetics of the synergistic effect, we continuously monitored the cells using a real-time apoptosis/necrosis assay. As shown in the right panel of Figure 1M, the apoptotic signal was already elevated 4h after the beginning of the combination treatment, while necrosis did not occur within the observed time frame.

To confirm these results in patient-derived material, we used HCC organoids described previously to be sorafenib-resistant [37]. Maintaining these tumoroids in a starvation-mimicking medium did not *per se* impede their viability (Figure 1N-O). However, compared to full expansion medium [38], starvation-mimicking medium elicited a dose-dependent decrease in viability in response to sorafenib (Figure 1N-O), resembling results in HepG2 cells (e.g. Figure 1J).

Opposing the sensitization of transformed and resistant HCC models, the viability of mouse primary hepatocytes was largely unaffected by sorafenib with or without starvation in different exposure protocols (Figure S1L-M). This suggests that sorafenib selectively targets HCC cells, while leaving non-transformed hepatocytes unharmed, which is mirrored by our pilot experiments in C57BL/6J mice (Figure S1G-H). Furthermore, our data are in line with a recent study showing beneficial effects on liver pathology of sorafenib at doses similar to the ones used in our study [39].

Altogether, these results show that nutrient restriction synergizes with sorafenib to alleviate therapy resistance by activating cell death programs in cultivated cells, HCC organoids, and xenografts.

### Sorafenib/starvation synergism is specific, but does not act via canonical kinase inhibition or single starvation-responsive pathways

To test if the synergism between starvation and sorafenib is a specific phenomenon, we next investigated co-treatment with starvation and doxorubicin, a chemotherapeutic that has been traditionally used in chemoembolization for intermediate-stage HCC [40]. Effective doses of doxorubicin (shown by p53 stabilization in response to doxorubicin-induced DNA damage in Figure S2A) led to a marginal decrease in cell viability under starvation conditions (Figure 2A). Consistent with previous clinical studies that showed no beneficial effects of doxorubicin over sorafenib treatment or after the failure of sorafenib treatment [41,42], we did not observe any additional effect over sorafenib/doxorubicin combination treatment, regardless of the cells being maintained in full growth medium or starvation medium (Figure 2B). Hence, these data suggest that the sensitization of sorafenib-resistant HCC cells is a specific effect conferred by the combination of sorafenib and starvation.

**Figure 2.**
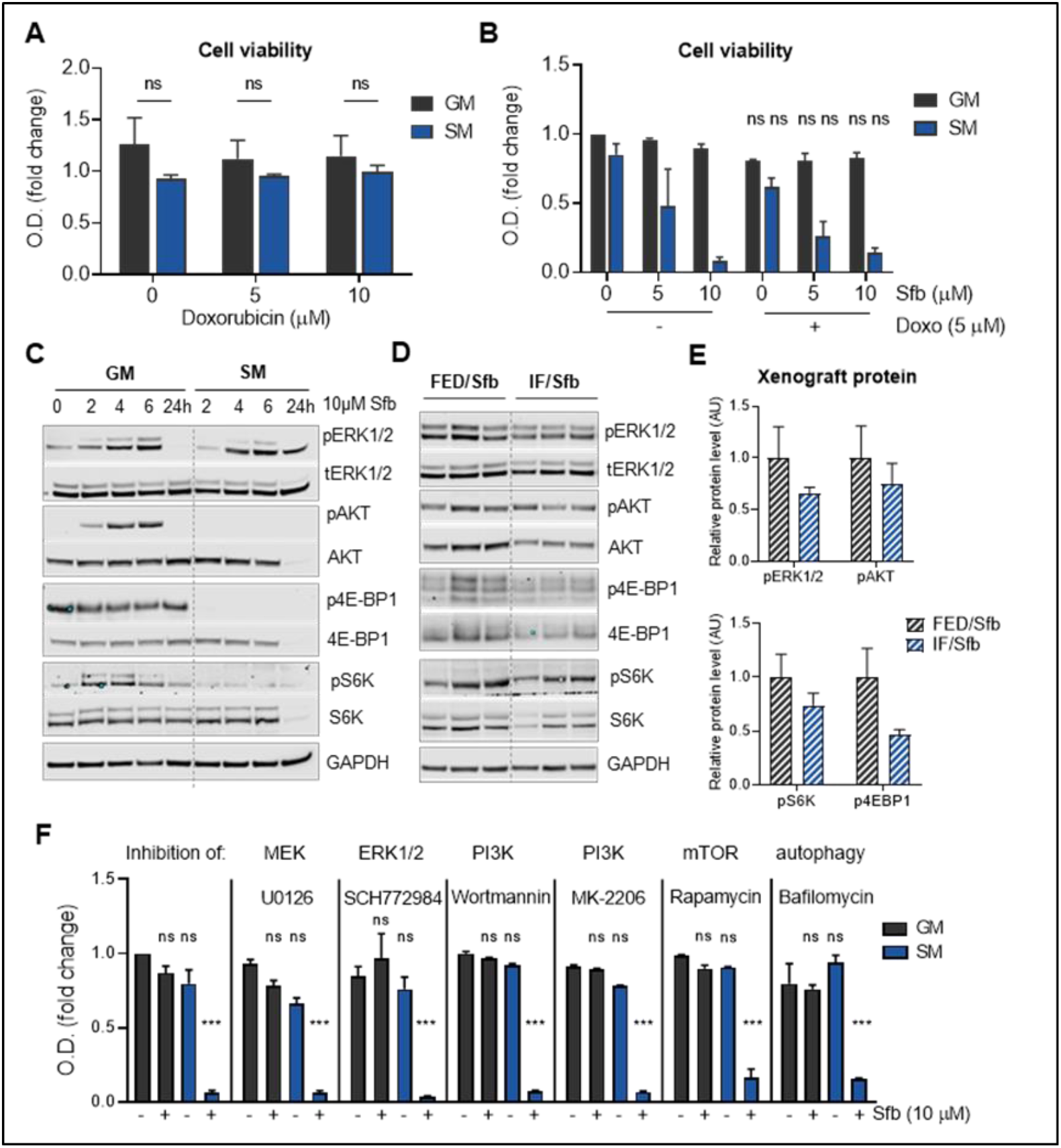
Sorafenib/starvation synergism is specific but does not act via canonical kinase inhibition or single starvation-responsive pathways. (A) Viability of HepG2 cells after 24 h of incubation in growth medium (GM) or starvation medium (SM) with indicated concentrations of doxorubicin (Doxo). Values are normalized to the viability of non-treated cells grown in GM. (B) Viability of HepG2 cells after 24 h of incubation in GM or SM with indicated concentrations of Doxo and/or sorafenib (Sfb). Values are normalized to the viability of non-treated cells grown in GM. Comparison of groups treated with Doxo and Sfb vs corresponding groups treated with Sfb is shown. (C) HepG2 cells were grown in GM or SM with 10 μM sorafenib for indicated times, and western blot analysis was performed to determine protein levels of AKT, ERK, 4E-BP1, S6K and their respective phosphorylated forms. GAPDH served as loading control. (D) Western blot analysis was performed to determine protein levels of AKT, ERK, 4E-BP1, S6K and their respective phosphorylated forms in HepG2 xenografts from ad libitum fed (FED) and intermittently fasted (IF) nude mice treated with 30 mg/kg Sfb. GAPDH served as loading control. (E) Quantification of phosphorylated protein levels, relative to GAPDH and normalized to the protein level in xenograft protein lysates from FED mice. Unpaired t-test was performed. (F) Viability of HepG2 cells after 24 h of incubation GM or SM with Sfb and/or MEK inhibitor (5 µM U0126), ERK1/2 inhibitor (1 µM SCH772984), PI3K inhibitor (1 µM Wortmannin), PI3K inhibitor (1 µM MK-2206), 10 nM Rapamycin and 10 nM Bafilomycin determined. Comparison of groups within each inhibitor treatment group are shown. If not noted otherwise, mean values ± SEM are shown and two-way ANOVA, Tukey’s multiple comparisons test was performed. *** p<0.001, ** p<0.01, * p<0.05, ns-not significant (p>0.05); O.D. optical density.

Sorafenib was initially identified as a RAF inhibitor in the RAS-RAF-MEK-ERK pathway [28]. Instead of inhibiting this pathway, sorafenib led to increased phosphorylation of ERK (Figure 2C) in resistant HepG2 cells, reminiscent of previous reports describing a paradoxical activation of this pathway by sorafenib and other BRAF inhibitors [43,44]. To test whether sorafenib acts via inhibition of other growth pathways, we investigated the fasting-responsive PI3K-AKT-mTOR pathway [13,45], hyperactivation of which is a common denominator of malignant reprogramming in many cancers [46,47]. Probing activation of AKT through phosphorylation of serine 473, we found that sorafenib treatment leads to a dynamic activation of AKT (Figure 2C). This could represent a compensatory response to boost glycolysis as reported in other cancer models [12] and a potential mechanism of resistance to sorafenib in our system. However, in starvation medium that only contains low glucose (5.5 mM, as opposed to 25 mM in full growth medium) and no growth factors, AKT activity is completely abrogated (Figure 2C), coinciding with sensitivity to sorafenib. mTORC1 was also activated under sorafenib treatment in normal growth medium but was completely blunted through starvation as evidenced by a loss of phosphorylation of mTORC1 substrates S6K1 and 4EBP-1 (Figure 2C). This reduction of growth pathways by combination treatment was largely recapitulated by analyzing tumors from the xenograft assay (Figure 2D-E).

Interestingly, targeted pharmacological inhibition (Figure S2B-E) of MEK, ERK, PI3K, AKT, mTORC1, as well as of autophagy (downstream of mTORC1)[47] did neither recapitulate sorafenib effects under starvation nor show any additional effects over starvation/sorafenib co-treatment on cell viability (Figure 2F). This set of experiments rules out these pathways as the sole mediators of starvation-mediated sensitization to sorafenib.

Thus, the paradoxical upregulation of several growth pathways in response to sorafenib treatment could constitute a resistance mechanism in our model [31]. Overcoming this resistance seems not to rely on single growth pathways but rather on the pleiotropic effects of starvation impacting on the PI3K-AKT-mTORC1 axis that, in combination with non-canonical sorafenib action, leads to HCC cell death.

### Sorafenib-induced Warburg shift is curtailed by glucose limitation under starvation

Several previous studies have described the impact of sorafenib on mitochondrial bioenergetics [39,48–50]. Furthermore, AKT activation boosts aerobic glycolysis in cancer cells with defective mitochondria [51]. We therefore investigated if such a Warburg shift, a hallmark of cellular transformation [12], could be the underlying mechanism of resistance in our model.

We first evaluated oxygen consumption rate (OCR), reflecting mitochondrial oxidative phosphorylation (OXPHOS), in sorafenib-resistant HepG2 cells after 6 hours of treatment, when cell viability was uncompromised in all conditions (Figure S3A) and AKT is maximally activated (Figure 2C). As shown in Figure 3A and B, starvation led to a dampening, and sorafenib treatment to a severe reduction, of the basal and stimulated OCR. Showing a potent additive effect, combination treatment completely obliterated any cellular respiration within 6 hours after treatment. Basal OCR, ATP-linked OCR, as well as maximal respiration were significantly reduced by either treatment alone and largely unmeasurable after the combination treatment (Figure 3B). Refeeding cells for 2 hours after pre-treatment with sorafenib, starvation, or both, showed that the starvation-mediated OCR attenuation is completely reversible while sorafenib treatment caused a prolonged inhibition of cellular respiration (Figure 3C), indicative of persistent inhibition of OXPHOS. Next, we determined the mitochondrial membrane potential. Starvation alone did not impact on mitochondrial membrane potential (Figure S3B). However, 2 and 6 hours of sorafenib treatment significantly reduced mitochondrial membrane potential (Figure 3D) while leaving mitochondrial morphology intact (Figure 3E). Because the effect of sorafenib on mitochondrial ATP-linked OXPHOS was most pronounced in growth medium (Figure 3A-B), we reasoned that the cells might execute a Warburg shift towards aerobic glycolysis to enable survival. Thus, we monitored glycolytic function in real-time upon acute treatment with sorafenib and oligomycin, an ATP-synthase inhibitor. Sorafenib was as potent as oligomycin in stimulating the extracellular acidification rate (ECAR), a proxy for glycolytic activity (Figure 3F). A six-hour sorafenib treatment in full growth medium led to increased levels of ECAR from non-glucose substrates (measurements before glucose injection), and after glucose and oligomycin addition, consistent with a Warburg shift (Figure 3G, left panel). However, cells pre-treated in starvation medium showed no metabolic flexibility upon sorafenib treatment (Figure 3G, right panel), even though acute addition of glucose can reinstate glycolysis to the maximal glycolytic capacity (Figure 3G, right panel). These data led us to hypothesize that, while sorafenib induces persistent OXPHOS inhibition eliciting a Warburg shift to glycolysis, starvation can acutely curtail this shift and thereby reduce cancer cell survival. This would imply that starvation utilizes a metabolic window of opportunity opened by sorafenib action as an OXPHOS inhibitor in our system. Of note, the starvation medium contains low levels of glucose (5.5 mM). Therefore, after 6 hours of treatment, this glucose level might still be sufficient for starved and sorafenib treated cells to remain viable (Figure S3A) by increasing their glycolysis rate (shown by increased ECAR in Figure 3B).

**Figure 3.**
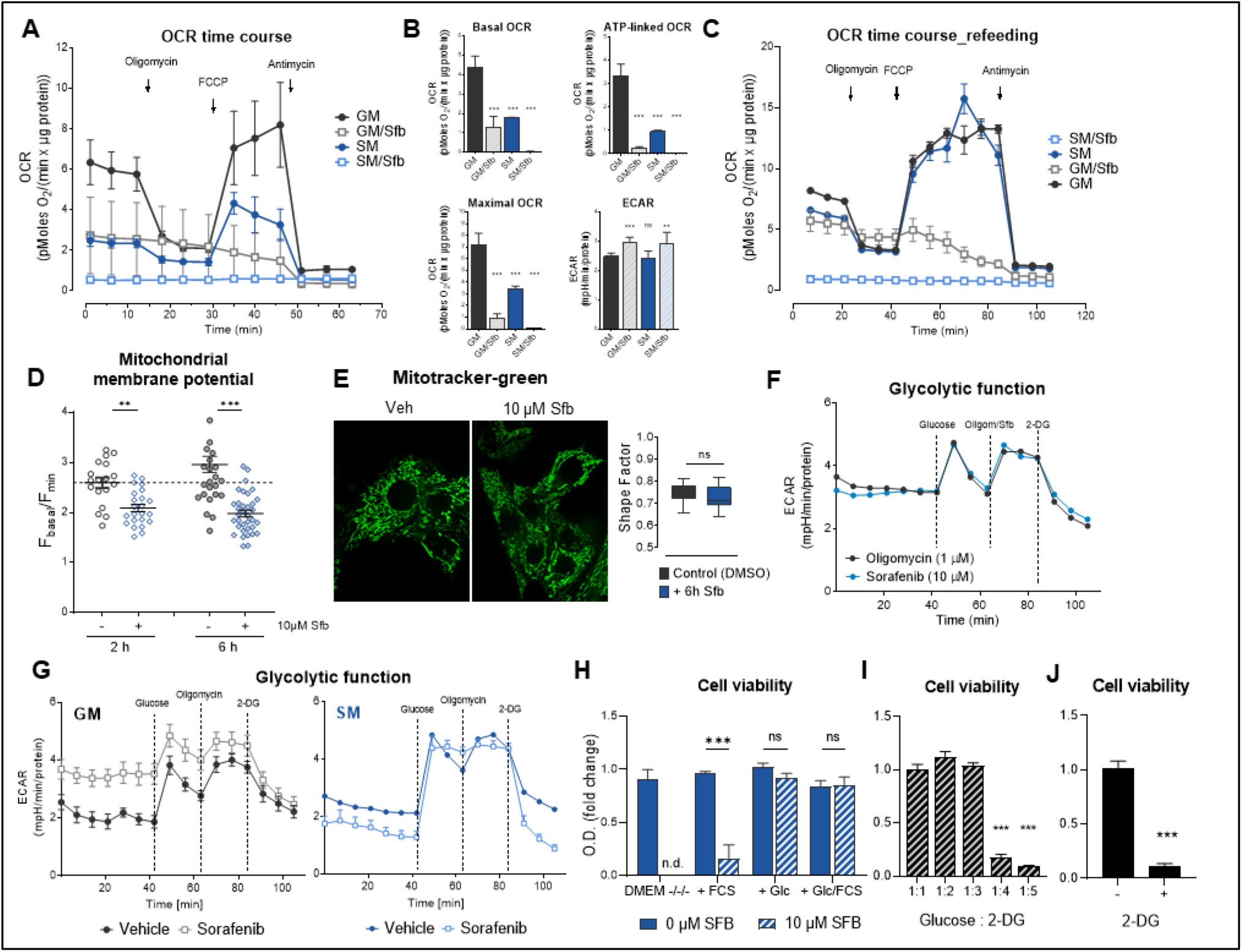
Sorafenib-induced Warburg shift is curtailed by glucose limitation under starvation. (A) Seahorse analysis of mitochondrial respiration in HepG2 cells. Cells were incubated 6 hours in growth medium (GM) or starvation medium (SM) with 0 and 10 μM sorafenib (Sfb). During the measurement, cells were kept under the same conditions and consecutively injected with oligomycin (4 μM), FCCP (0.2 μM), and antimycin (2.5 μM). Continuous oxygen consumption rates (OCR) values are shown. (B) Basal respiration, ATP-linked respiration, maximal respiration, and extracellular acidification rate (ECAR) in four treatment conditions. Comparison to GM group is shown. (C) Seahorse analysis of mitochondrial respiration in HepG2 cells. Cells were incubated for 6 hours in GM or SM with 0 and 10 μM Sfb. Then, the media were replaced by fresh GM and cells were incubated for an additional 2 hours. Measurements were performed as described I (A). Continuous OCR values are shown. (D) Mitochondrial membrane potential measured using TMRM in HepG2 cells after 2- or 6-hours incubation with vehicle (DMSO) or 10 µM Sfb. Unpaired t-test was performed. (E) Mitotracker green mitochondrial staining in HepG2 cells (left panel) after 6 hours treatment with 10 µM Sfb and signal quantification (right panel). Unpaired t-test was performed. (F) Glycolysis stress test in HepG2 cells grown in GM. For the measurements, cells were incubated in glucose-dree XF assay medium supplemented with 2 mM glutamine and consecutively injected with glucose (10 mM), oligomycin (1 μM), or Sfb (10 µM) and 2-deoxyglucose (50 mM). Continuous ECAR values are shown. (G) Confluent HepG2 cells were grown in GM (left panel) or SM (right panel) with or without 10 µM Sfb for 6 hours before glycolytic function assay was performed. Continuous ECAR values are shown. (H) Viability of HepG2 cells after 24 hours incubation in DMEM without glucose, glutamine, and pyruvate (DMEM) or with added 10% FCS, 25mM glucose, or both, with or without 10 µM Sfb. (I) HepG2 cells were kept in GM supplemented with 10 µM Sfb and with the addition of 2-Deoxy-D-glucose (2-DG) in indicated molar ratios to glucose. After 24 hours, a viability assay was performed. One-way ANOVA, Tukey’s multiple comparisons test was performed, all groups compared to 1:1 group. Y-axis same as in 3H. (J) Organoid viability after 6 days of incubation in GM with 5 µM sorafenib and with or without 50 mM 2-DG (1:2 ratio to glucose). Unpaired t-test was performed. Y-axis same as in 3H. If not noted otherwise, mean values ± SEM are shown and two-way ANOVA, Tukey’s multiple comparisons test was performed. *** p<0.001, ** p<0.01, * p<0.05, ns-not significant (p>0.05); O.D. optical density.

To identify the major metabolite responsible for the sensitization effect of starvation we used DMEM without glucose, pyruvate, glutamine, and FCS (DMEM^-/-/-^) and observed the viability upon individual add-back of these components. Interestingly, and corroborating data from above showing that the addition of glucose immediately rescues glycolytic function (Figure 3G), only glucose add-back maintained cell viability (Figure 3H). Neither add-back of glutamine, nor pyruvate, rescued viability (Figure S3C). FCS, rich in growth factors, hormones, and metabolites, had a marginal effect in increasing viability upon addition to DMEM^-/-/-^(Figure 3H). This experiment indicated that glucose may be the primary metabolite responsible for the sorafenib sensitization effect of starvation. To directly test this, we treated cells kept in full growth medium and treated with sorafenib with increasing concentrations of the glycolysis inhibitor 2-deoxy-D-glucose (2-DG). Starting with a stoichiometric ratio of 1:4 (glucose:2-DG), cell viability was significantly decreased after 24 hours (Figure 3I), robustly establishing glucose as the limiting metabolite required for sorafenib-sensitization through starvation. This was confirmed for sorafenib-resistant HCC organoids (Figure 1M-N), as inhibition of glycolysis by 2-DG in full medium significantly reduced their viability (Figure 3J).

Altogether, our data indicate that sorafenib acts non-canonically as an OXPHOS inhibitor initiating a Warburg shift to aerobic glycolysis in HepG2 cells. Starvation or IF curtails this metabolic flexibility through the limitation of glucose and, with that, confers sensitivity towards the anti-cancer effects of sorafenib.

### p53 is required for synergistic anti-tumorigenic action of starvation and sorafenib

Loss-of-function of the tumor suppressor and transcription factor p53 is causative for or involved in the development of over 50% of human cancers [52] and ∼30% of HCC cases present with a mutation in the p53 coding gene [53]. Apart from its tumor suppressor function, p53 is increasingly appreciated as a regulator of cancer metabolism [54] and we recently published that the p53 protein is stabilized in starved HepG2 cells, leading to activation of downstream target genes [36].

To investigate the influence of p53 on starvation-mediated sensitization to sorafenib we used CRISPR/Cas9 to derive p53 knock-out (p53KO) clones from resistant HepG2 cells (Figure S4A-B). Interestingly, isogenic p53KO cells treated with the same starvation/sorafenib protocol used for their p53-proficient counterparts (Figure 1I) did not show a significant decrease in viability (Figure 4A) or an increase of cytotoxicity (Figure 4B), which renders p53-deficient cells resistant to sorafenib even under starvation conditions. To exclude potential clonal effects, we validated this observation in other CRISPR/Cas9 clones (Figure S4C) and re-expressed p53 in the p53KO cells (Figure S4D). The re-expression of p53 reinstated target gene expression (Figure S4E) and sensitization to sorafenib under starvation (Figure 4C, right panel), indicating that p53 is necessary and sufficient for this synergistic effect.

**Figure 4.**
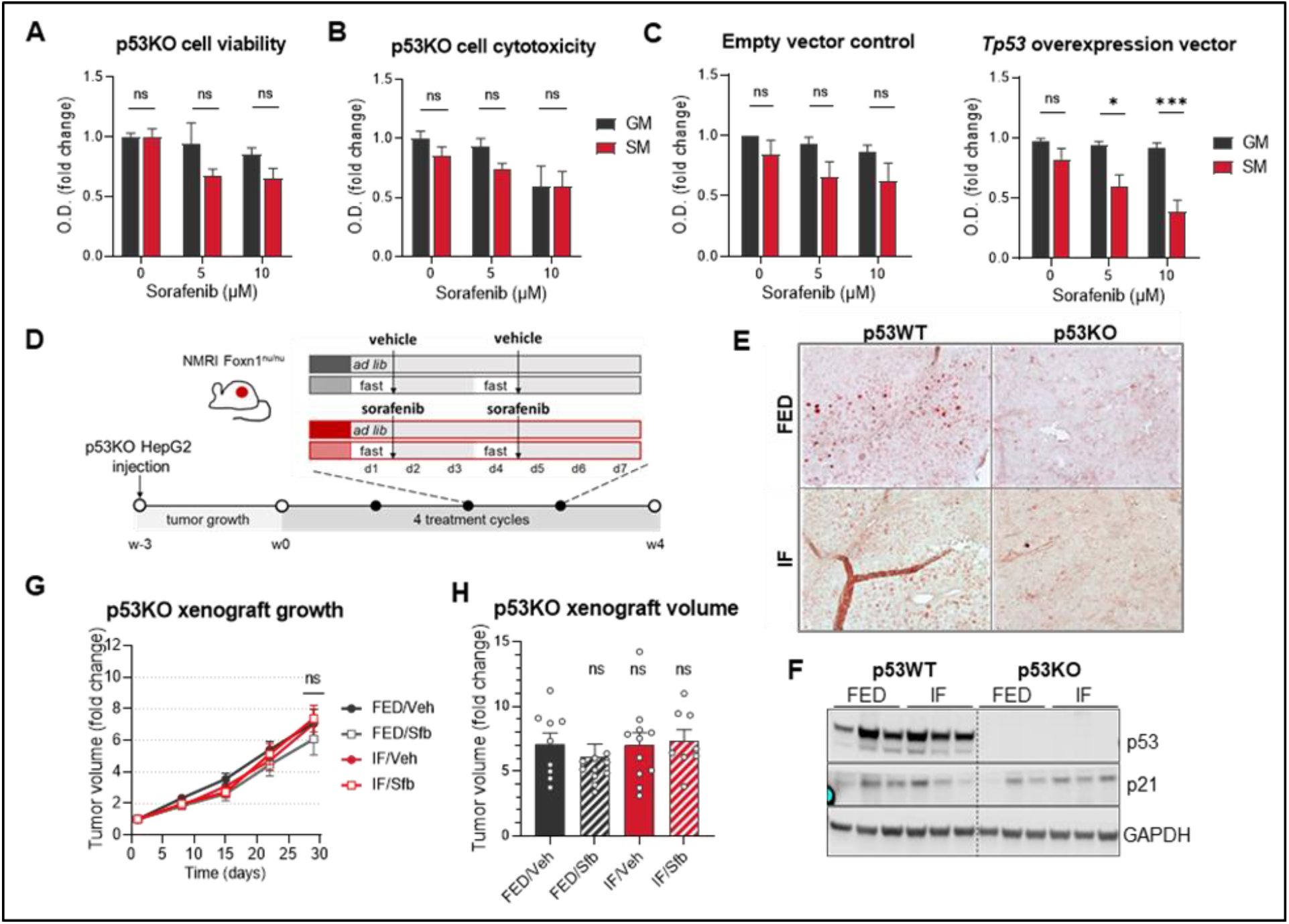
p53 is required for synergistic anti-tumorigenic action of starvation and sorafenib. (A) Viability of HepG2 p53KO cells after 24 hours of incubation in growth medium (GM) or starvation medium (SM) with indicated concentrations of Sfb. Values are normalized to the viability of non-treated cells grown in GM. (B) Cytotoxicity measured by LDH assay in p53KO cells grown in GM vs cells grown in SM with indicated Sfb concentrations after 24 hours. (C) p53KO cells were transfected with an empty vector (EV, left panel) and p53 overexpression vector (OE, right panel). Viability assay was performed after 24 hours of incubation in GM or SM with indicated concentrations of Sfb. (D) Experimental design: after p53KO HepG2 cells that were injected into hind flanks of NMRI Foxn1nu/nu mice formed palpable xenografts, mice were divided into four groups as described in figure 1A. (E) Immuno-histochemical staining of p53 in xenograft sections from mice injected with p53WT and p53KO HepG2 cells. (F) Western blot analysis was performed to determine protein levels of p53 and p21 in xenografts from ad libitum fed (FED) and intermittently fasted (IF) nude mice. GAPDH served as loading control. (G) Xenograft volume normalized to its starting volume; n=8-11 per group. Two-way ANOVA, Dunnett’s multiple comparisons test, all groups compared to FED/Veh group. (H) Relative xenograft volumes at day 29. One-way ANOVA, Tukey’s multiple comparisons test, comparison of each group vs control, FED/Veh group is shown; n=8-11 per group. If not noted otherwise, mean values ± SEM are shown and two-way ANOVA, Tukey’s multiple comparisons test was performed. *** p<0.001, ** p<0.01, * p<0.05, ns-not significant (p>0.05); O.D. optical density.

We next xenografted p53KO cells in nude mice and subsequently subjected them to the same IF/sorafenib protocol (illustrated in Figure 4D and S4F) as for p53-proficient HepG2 cells (p53 wild-type, p53WT) (Figure 1A). Immunohistochemistry (Figure 4E), western blot (Figure 4F), and qPCR (Figure S4G) assessing p53 and p53 target gene expression confirmed persistent knock-out throughout the *in vivo* tumor growth assay. Importantly, subcutaneous tumor growth rates were similar in all four groups (Figure 4G-H), confirming *in vivo* that p53 is required for the synergistic effect of IF/starvation and sorafenib.

### p53 confers starvation/sorafenib synergism by regulating glycolytic rate and capacity

To decipher the mechanistic aspects of p53 status on the synergism of sorafenib and starvation, we first investigated growth pathways and cancer cell metabolism. p53WT (Figure 2C) and p53KO cells, showed similar activation of ERK, AKT, and mTORC1 pathways when treated with sorafenib in growth medium and a similar abrogation of AKT and mTORC1 signaling in starvation medium (Figure 5A). Combination treatment completely blocked OXPHOS in p53KO just as in p53WT cells (Figure S5A), while the glycolytic rate and capacity in p53KO cells were lower (Figure 5B-C) when compared to p53WT cells. p53WT cells retain their glucose uptake at a similar level in both control and combination treatment, while p53KO cells significantly reduce glucose uptake upon combination treatment (Figure 5D-E). Corresponding with reduced glucose uptake, the most abundant glucose transporter in liver cells (*SLC2A2*) was strongly downregulated in p53-deficient cells (Figure 5F). Furthermore, glucose transporter downregulation (*SLC2A1, SLC2A2, SLC2A3*) was also observed in p53KO xenografts after four weeks of combined treatment (Figure 5G). Complete depletion of glucose severely dampened p53KO cells’ survival under sorafenib treatment and only add-back of glucose, but not pyruvate, glutamine, or FCS (Figure S5B), was sufficient to reinstate sorafenib resistance in p53KO cells (Figure 5H and S5B). Accordingly, blockage of glycolysis with 2-DG and concomitant sorafenib treatment led to p53KO cell death (Figure 5I). These results demonstrate that p53KO cells under combination treatment reduce the expression of glucose transporters, resulting in reduced glucose uptake and lower glycolytic rate and capacity compared to p53WT cells. While p53KO cells are as dependent on glucose as p53WT cells, their “economic” use of the little glucose in the starvation medium could prolong their survival.

**Figure 5.**
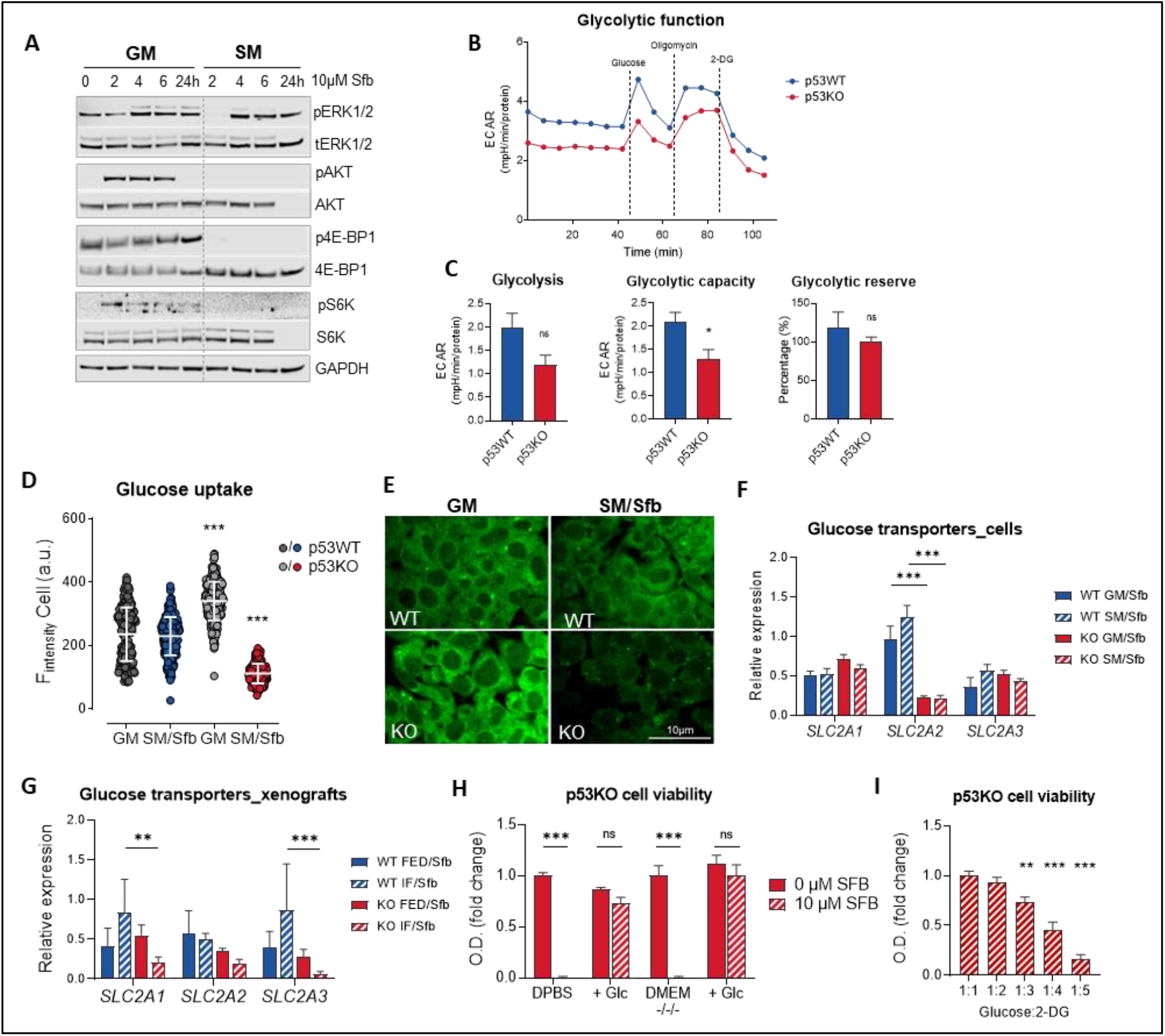
p53 confers starvation/sorafenib synergism by regulating the glycolytic rate. (A) p53KO HepG2 cells were grown in growth medium (GM) or starvation medium (SM) with 10 μM sorafenib (Sfb) for indicated times, and western blot analysis was performed to determine protein levels of AKT, ERK, 4E-BP1, S6K and their respective phosphorylated forms. GAPDH served as loading control. (B) (C) Glycolysis stress test (GST) in HepG2 p53WT and p53KO cells. Cells were grown in GM for 24 hours before GST was performed. For the measurements, cells were incubated in glucose-dree XF assay medium supplemented with 2 mM glutamine and consecutively injected with glucose (10 mM), oligomycin (1 μM) or Sfb (10 µM), and 2-deoxyglucose (50 mM). Unpaired t-test was performed. (D) Glucose uptake measurement. p53WT and p53KO cells were kept in GM or SM with 10 µM Sfb for 6 hours before glucose uptake was analyzed. One-way ANOVA, Tukey’s multiple comparisons test was performed. Comparison of all groups vs. GM is shown. (E) Representative images from the glucose uptake assay. (F) Expression of *SLC2A1, SLC2A2*, and *SLC2A3* determined in HepG2 WT and KO cells after 24-hour incubation. Relative to reference genes *PPIA* and *B2m*. (G) Expression of *SLC2A1* and *SLC2A3* determined in HepG2 WT and KO cells xenografts after four weeks of treatment. Relative to reference genes *PPIA* and *B2m*. (H) p53KO HepG2 cells were kept in DPBS or DMEM^-/-/-^ alone or with the addition of 25 mM glucose with or without 10 µM Sfb. After 24 hours, a viability assay was performed. (I) p53KO HepG2 cells were kept in GM supplemented with 10 µM Sfb and with the addition of 2-DG in indicated molar ratios to glucose. After 24 hours, a viability assay was performed. One-way ANOVA, Tukey’s multiple comparisons test was performed. Comparison of all groups vs. 1:1 group is shown. If not noted otherwise, mean values ± SEM are shown and two-way ANOVA, Tukey’s multiple comparisons test was performed. *** p<0.001, ** p<0.01, * p<0.05, ns-not significant (p>0.05); O.D. optical density.

### p53 coordinates the pro-apoptotic proteome during combination treatment

To further dissect why p53 is necessary for the synergistic action of fasting/starvation and sorafenib we performed proteomics comparing p53WT and p53KO samples from cells and xenografts treated with sorafenib and starvation or sorafenib and IF, respectively. Overall, 5064 proteins were reproducibly detected and quantified in all triplicate experiments with the p53WT and p53KO cells and the concomitant p53WT and p53KO xenografts. Of these, 2770 proteins were significantly changed, as determined by using ANOVA testing (FDR 0.05). Hierarchical clustering correctly assigned replicates to experimental groups and identified four clearly distinct clusters of expression profiles (Figure 6A). While clusters 1 and 4 are largely comprised of proteins with varying expression levels in cells and xenografts, clusters 2 and 3 show proteins whose expression profiles were influenced in a similar way by p53KO (Figure 6A). As the activity of p53 has been shown to be confined to transcriptional activation [55], we focused on cluster 3 containing 147 proteins downregulated in p53KO samples (Figure 6B). To further distinguish between direct and indirect effects of p53 knock-out, we overlapped these 147 proteins with high-confidence p53 targets derived from a previous meta-analysis [56]. From this approach, only eight p53 targets emerged in our proteomics data set as consistently downregulated through p53KO both *in vivo* and *in vitro* (Figure 6C and 6D). Interestingly and consistent with increased apoptosis upon combination treatment (Figure 1L and 1M), the pro-apoptotic regulator BAX was among the eight candidates (Figure 6D), which was confirmed on protein and mRNA level for cells (Figure 6E and 6F) and xenografts (Figure 6G and 6H).

**Figure 6.**
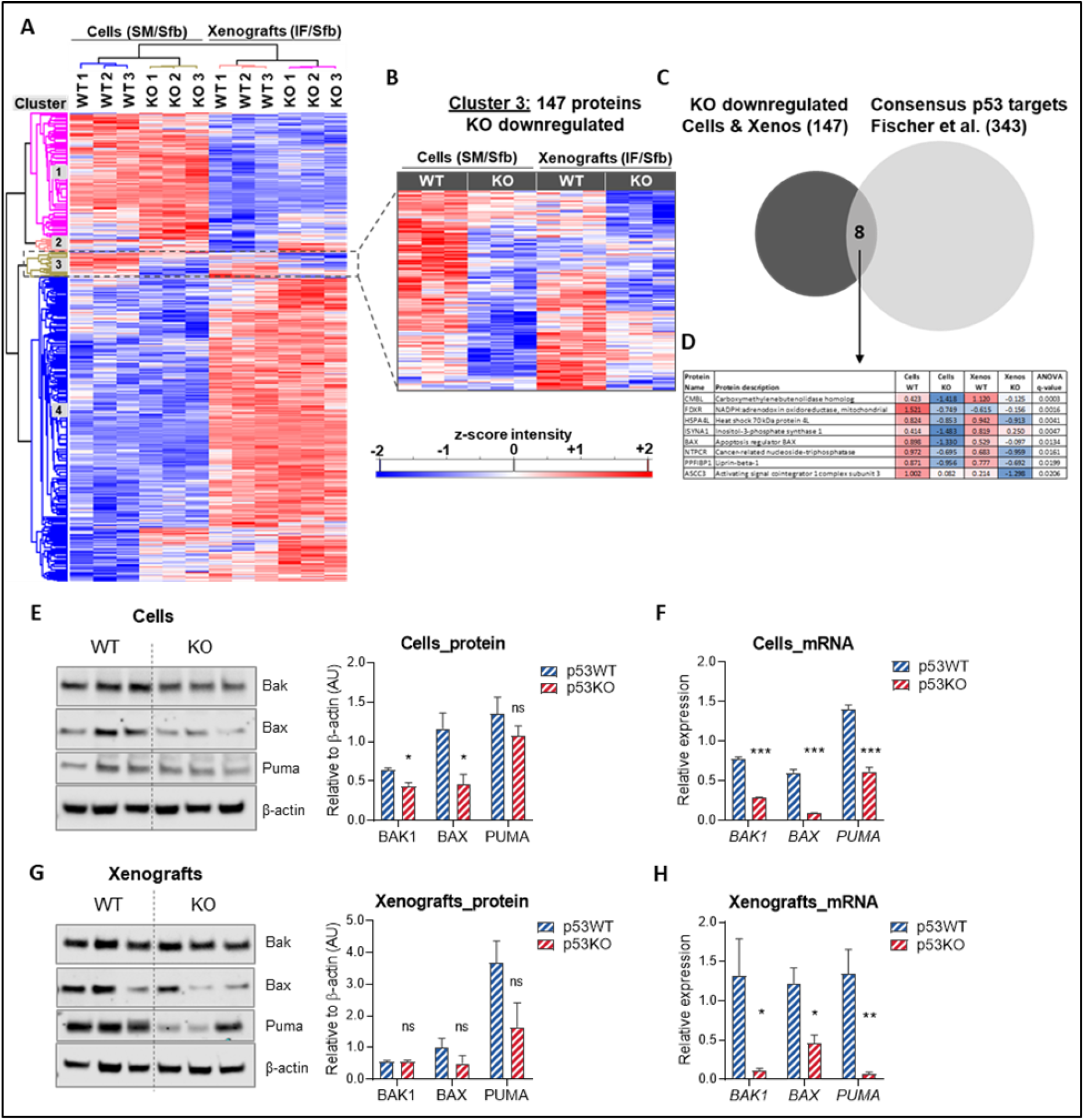
p53 regulates pro-apoptotic proteome under starvation/sorafenib combination treatment. (A) Heatmap of proteomics analysis from lysates derived from p53WT and p53KO HepG2 cells or xenografts. Xenografts were treated with IF and sorafenib as described in Figure 1A and 4D. HepG2 cells were grown in starvation medium (SM) with 10 μM sorafenib (Sfb) for 6 hours. Proteins detected in all samples were submitted to ANOVA testing (FDR 0.05) and to hierarchical clustering using Perseus. Values are displayed as z-score. (B) Heatmap of 147 proteins from cluster 3. Values are displayed as z-score. (C) Cluster 3 proteins were overlapped with biochemically verified p53 target genes [56]. (D) Eight proteins from overlap are shown with averaged z-scores and ANOVA q-values. (E) Western blot analysis and densitometric quantification measuring BAX, BAK and PUMA protein levels from p53WT and p53KO cells after 6-hour treatment with 10uM sorafenib in SM. Samples from three independent experiments were used. β-actin served as loading control. (F) qPCR expression analysis for samples as in (E). Relative to reference genes *PPIA* and *B2M*. (G) Western blot analysis and densitometric quantification measuring BAX, BAK, and PUMA protein levels from p53WT and p53KO xenografts after 4 weeks of IF/sorafenib combination treatment. Samples from three different xenografts with the same treatment were used. β-actin served as loading control. (H) qPCR expression analysis for samples as in (E). Relative to reference genes *PPIA* and *B2M*. If not noted otherwise, mean values ± SEM are shown and Unpaired t-test was performed. *** p<0.001, ** p<0.01, * p<0.05, ns-not significant (p>0.05).

We next established that two other pro-apoptotic proteins, BAK1 and PUMA, described to be regulated by p53 [52,57] but not detected in the proteomics analysis, were downregulated in p53KO groups (Figure 6E-H). These data suggest that p53 is essential to prime HCC cells for apoptosis, resulting in higher susceptibility to intrinsic cell death upon combination treatment.

### Intermittent fasting improves sorafenib action in an orthotopic HCC mouse model

To investigate sorafenib/starvation synergism beyond the alleviation of resistance in late-stage HCC, we used a sorafenib-responsive HCC-derived cell line. Huh-6 clone 5 cell line is *per se* responsive to sorafenib in a dose-dependent manner (Figure S6A). Simultaneous starvation significantly improved sorafenib efficacy (Figure S6A), implying a synergistic anti-cancer effect in non-resistant HCC cells.

To test if this sorafenib-enhancing effect of starvation can be recapitulated in an orthotopic HCC mouse model, we treated mice with a single dose of diethylnitrosamine (DEN), a well-established protocol to induce HCC nodules in livers within months [58]. The mice used were transgenic for a liver-specific conditional KO of p53 (AlbCreER^T2^ X p53^lox/lox^ mice [59] (Figure S6B – S6E)). In this model, p53KO was successfully established (Figure 7A) through oral gavage of 100 mg/kg of tamoxifen after liver nodules developed (ultrasound imaging, Figure S6F) and before the treatment protocol was started (Figure 7B). This experimental timeline was important to dissociate the impact of p53 status on treatment effects from long-term effects of p53 deficiency on tumorigenesis, as would be expected in a germ line p53KO model [60]. After four weeks of treatment (ad libitum fed (FED) + sorafenib or IF + sorafenib), total body weight measured after refeeding phases (Figure 7C) was unchanged. In addition to terminal liver weight (Figure 7D), we found that in the p53-proficient HCC nodule weight was reduced in the IF/sorafenib group as compared to the FED/sorafenib group (Figure 7E, upper panel and whole liver photographs in lower panel). However, in the p53KO group, this IF-mediated reduction was absent, indicating that p53 is necessary for sorafenib-potentiating effects of IF (Figure 7E), reminiscent of the situation in p53KO HepG2 xenografts (Figure 4G-H). Combination treatment also decreased nodule numbers in p53-proficient livers of DEN-treated mice (Figure S6G). In line with the changes in HCC nodule weight, concentrations of plasma markers of liver injury such as AST, ALT, and LDH were notably lower in the p53WT group after combination treatment when compared to the p53KO group (Figure 7F). However, other liver markers remained unchanged (Figure S6H). Furthermore, the expression of glucose transporters 1 and 2 was decreased in p53KO HCC nodules in the combination treatment group in comparison to nodules in p53-proficient livers (Figure 7G). This result is in line with the reduced glucose uptake shown in p53KO HepG2 cells (Figure 5D-E) and the reduction of glucose transporter expression in p53KO xenografts (Figure 5G). The pro-apoptotic genes *Bak1, Bax*, and *Puma* were downregulated in the p53KO groups and/or by combination treatment (Figure 7H). Collectively, our data from non-resistant HCC cell lines and an orthotopic HCC mouse model indicate a therapy-enhancing effect of starvation/IF in combination with sorafenib, warranting further studies to determine whether sorafenib can be utilized in early-stage HCC when combined with IF regimens.

**Figure 7.**
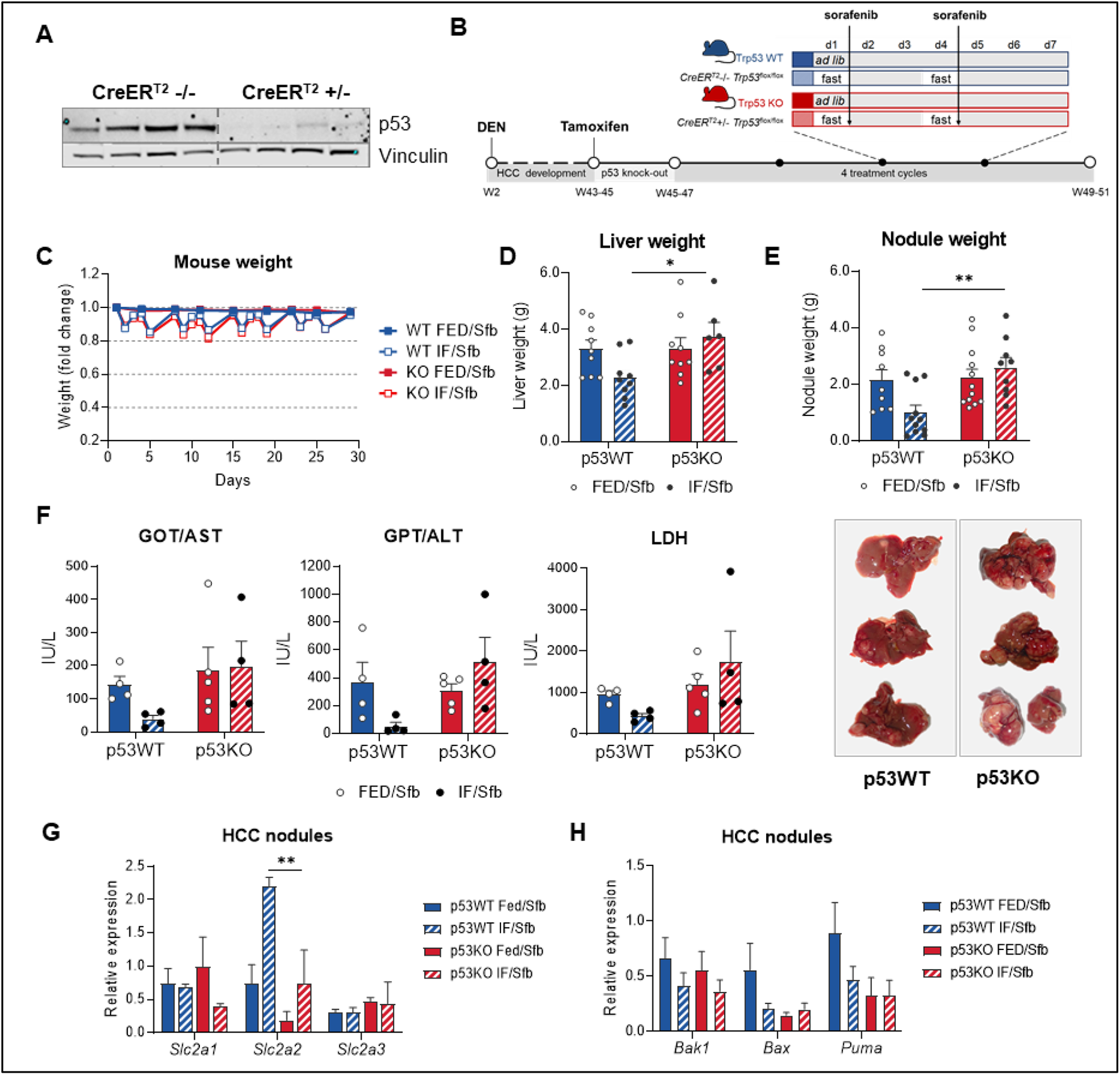
Intermittent fasting improves sorafenib action in an orthotopic HCC mouse model. (A) Western blot was performed with the protein lysates from HCC samples after the treatment to assess p53KO in the livers. Vinculin served as loading control. (B) CreER^T2+/-^and CreER^T2-/-^Tp53^flox/flox^ mice received 25 mg/kg N-Nitrosodiethylamine (DEN) i.p. at the age of two weeks. After 41-43 weeks, after developing HCC, the mice received 100 mg/kg tamoxifen by oral gavage for five consecutive days, followed by one week wash-out phase. Then, 4 cycles of treatment were conducted: mice were either fed ad libitum (FED) or fasted two times per week for 24 hours in the intermittently fasted (IF) group. Both groups received 30 mg/kg Sfb two times per week, after the 24-hour fasting in the IF group, and on the same day in the FED group. After 4 cycles, and additional 2-3 days of ad libitum feeding, the mice were sacrificed. (C) Animal weight during the 4-week protocol. (D) Liver weight relative to body weight at the end of the experiment. (E) HCC nodule weight relative to body weight at the end of the experiment (upper panel). Photographs of livers from the IF/Sfb groups at the end of the experiment (lower panel). (F) Concentrations of ALT, AST, and LDH in the plasma on the sacrifice day in mice from designated groups. (G) Expression of glucose transporter genes determined in HCC nodules from FED/Sfb and IF/Sfb groups. Relative to reference genes *Ppia* and *B2m* (H)Expression of pro-apoptotic genes determined in HCC nodules from FED/Sfb and IF/Sfb groups. Relative to reference genes *Ppiaß* and *B2m* If not noted otherwise, mean values ± SEM are shown and two-way ANOVA, Tukey’s multiple comparisons test was performed. *** p<0.001, ** p<0.01, * p<0.05, ns-not significant (p>0.05).

## DISCUSSION

Overall, our data provide evidence that nutrient restriction can alleviate sorafenib resistance in human HCC models and that it can improve sorafenib efficacy in non-resistant settings. Our data establish a metabolic synergy between sorafenib-mediated OXPHOS inhibition and glucose limitation by nutrient restriction, imposing a “double-whammy” on cancer cell metabolism (see graphical abstract). This observation, if generalizable, creates a range of novel combination therapy opportunities for HCC treatment: the combination of glucose limiting regimens, or anti-glycolytic agents, with OXPHOS inhibitors. In support of this principle, previous studies have shown that suppression of glycolysis through targeting Hexokinase-II [61,62], or LDH [63], increased sorafenib efficacy consistent with our results in non-resistant models (Figure 7). In Huh-7 cells, combined treatment with sorafenib and the pyruvate dehydrogenase kinase inhibitor DCA [64] or aspirin (which inhibits glycolysis via PFKPFB3 inhibition)[65] reversed acquired sorafenib-resistance. Such attempts have been shown to be successful in other cancer entities as well. For instance, the combination of glucose restriction and metformin (shown to act as a mitochondrial complex I inhibitor) [66], led to restriction of tumor growth in mice xenografted with colon cancer and melanoma cells [67]. Collectively, these examples, together with our results, indicate that targeting mitochondrial metabolism [68] can be successfully combined with strategies that curb glycolysis [69]. However, pharmacologic targeting of glycolysis might elicit adverse effects on non-transformed cells [68] or provoke the emergence of cancer cells that develop resistance by switching to utilization of other metabolites or nutrients [70]. The pleiotropic and systemic effects of fasting on growth pathways and availability of various metabolite substrates [71,72], on the other hand, could squelch alternative tumorigenic pathways while being safe for non-transformed cells that have evolved coping with frequent fasting periods [8]. In fact, the first clinical trials that evaluate fasting as an adjuvant to cancer therapy attest a positive safety profile [8,73].

In line with our data on p53, a more recent study in melanoma and breast cancer cells showed the Rev1-p53 axis as necessary for the anti-tumorigenic effects of starvation through regulating the expression of pro-apoptotic genes during combined treatment [74]. Our data in HCC cells suggest a similar priming to intrinsic cell death [75,76] in cells proficient for p53, whereas p53 loss led to reduced expression of pro-apoptotic players. Especially, the pro-apoptotic effector Bax, shown to be a direct transcriptional target of p53 [77], was consistently downregulated *in vitro* as well as *in vivo*, possibly allowing p53KO cells to resist apoptotic stimuli through the relatively short starvation periods in our starvation/IF protocols. Since BH3 mimetics have been shown to sensitize HCC-derived cells to sorafenib [78], this might be a useful additional therapy to overcome the reinstated resistance in p53KO entities.

In addition to p53-mediated apoptosis, we found downregulation of several glucose transporters in the HepG2 xenografts and HCC nodules of DEN-treated mice in p53-deficient versus p53-proficient samples. Although reporter assays showed a repressive effect of p53 on *GLUT1* (*SLC2A1*) and *GLUT4* (*SLC2A4*) promoters [79], p53 may be necessary for regulation of glucose transporters in the chromatin context or in a cellular context that might involve a certain set of p53 effectors, including miRNAs [80]. However, the requirement of p53 for an effective Warburg shift in HCC cells is in agreement with p53’s role as tumor suppressor, coordinating not only cell cycle progression and apoptosis but in particular cancer cell metabolism [54,81,82], in some instances not complying with the standard notion of its function on glycolysis and OXPHOS regulation [83].

Taken together, the data presented herein could open new therapeutic windows to improve late-stage HCC therapy and should inspire clinical trials that apply fasting/sorafenib combination therapy, even for early/mid-stage HCC where currently no molecular therapy is efficacious. Furthermore, our data on p53 is clinically meaningful as it suggests to include a patient’s p53 status in the diagnostic panel before embarking on such combination therapy.

## MATERIALS AND METHODS

### Cell culture

HepG2, Hep3B, and Huh6 clone5 cell lines were purchased from ATCC and were cultivated in growth medium (GM) consisting of Dulbecco’s Modified Eagle Medium (DMEM, Gibco, Life Technologies) containing 25mM (4.5 g/L) D-Glucose, supplemented with 4 mM L-Glutamine (Glutamax™, Gibco, Life Technologies), 10% fetal calf serum (FCS, GE Healthcare Life Sciences) and 500 U/mL penicillin-streptomycin (Gibco, Life Technologies) at 37°C in humidified atmosphere with 5% CO_2_. Starvation medium (SM) consisted of Hank’s Balanced Salt Solution (HBSS) with CaCl_2_ and MgCl_2_ (Gibco, Life Technologies) containing 5.5 mM (1g/L) D-Glucose, supplemented with 10 mM HEPES (Gibco, Life Technologies).

Following cell culture supplements were used: 2-Deoxy-D-glucose (2DG) (Santa Cruz Biotechnology), D-Glucose (Sigma Aldrich), Sodium Pyruvate (Gibco, life Technologies), Dulbecco’s Phosphate-Buffered Saline (PBS, Gibco, Life Technologies), DMEM w/o D-glucose, L-glutamine and sodium pyruvate (Gibco, Life Technologies). Following therapeutics were used: Sorafenib (LC Laboratories) was dissolved in DMSO to 10 mM stock solution for cell culture use, Doxorubicin (Pfizer).

Following inhibitors were used: SCH 772984 ERK1/2 inhibitor (Cayman Chemicals), U0126 MEK inhibitor (Promega), Wortmannin (HY-10197-10mM/1mL), Buparlisib (HY-70063-10mM/1mL) and MK-2206 dihydrochloride (HY-10358-10mM/1mL) all from MedChemExpress, Bafilomycin A1 (BioMol), Rapamycin (Life Technologies, NOVEX).

### Primary mouse hepatocytes

C57BL/6J mice were anesthetized with 4 µl/g Ketamin (Pfizer)/Xylazin (Bayer) (8% to 1.2%) mixture. Liver was perfused by pumping (4 ml/min for 5 min) a pre-warmed perfusion buffer (EBSS w/o CaCl_2_/MgCl_2_ with 0.5 mM EGTA, Gibco, Life Technologies) through a catheter inserted into the *vena cava*. Liver was then digested by a pre-warmed digestion buffer consisting of HBSS and 5000 U collagenase I (Worthington Biochemical). The liver was then excised and cut on ice in GM and passed through a cell strainer. The cell suspension was centrifuged at 770 rpm for 3 min to obtain a cell pellet. Afterwards the pellet was resuspended in 25 ml of GM and 25ml of Percoll buffer (Biochrom) (containing 5 ml 10xPBS and 45 ml Percoll) and centrifuged at 770 rpm for 10 min. The supernatant containing dead cells was discarded and the pellet was again resuspended in 25 ml GM and centrifuged. 2×10^5^ cells were seeded on collagen-I (Corning) coated plates in GM.

### Patient-derived organoid culture

Patient HCC-derived organoids (the sorafenib-resistant HCC-1 line) [37] were kindly provided by Dr. Meritxell Huch (Max Planck Institute, Dresden, Germany). Organoids were thawed and expanded as previously described [38]. For experiments, organoids were harvested from two fully dense 50 µl Matrigel droplets (two wells from a 24-well plate) and seeded in 5 µl droplets per well in a 96-well plate, with 50 µl of media. GM was the same as organoid expansion medium [38] and SM comprised 1/3 DMEM w/o D-glucose, L-glutamine, and sodium pyruvate and 2/3 organoid expansion medium. Organoids were kept in GM or SM with or w/o sorafenib and/or 2DG for 6 days, with media and sorafenib replenishment every two days. Then, a viability assay was performed.

### Viability assay

For the viability assay, 2×10^5^ cells were seeded per well in a 96-well plate (Thermo Fisher Scientific) in 200 µL of GM to reach full confluency. Cells were allowed to attach for 24 h at 37°C and 5% CO_2_. After that, the medium was discarded, and cells were washed with PBS. Next, cells were treated as described in Results. Cell viability was analyzed using EZ4U assay (Biomedica Immunoassays) according to manufacturer’s instructions. Briefly, at the end of treatment the media was replaced with fresh GM (200 μl/well) and 20 μl/well EZ4U working solution. After 2 h of incubation at 37°C, the absorbance was measured at 492 nm with a reference-wavelength of 620 nm (Spark^®^ 10M multimode microplate reader).

### Gentian violet staining

The medium was discarded and the cells were stained by adding Carbol gentian violet solution (Roth). After 15 min of incubation, cells were washed with deionized water several times, until the dye stopped coming off. The cells were allowed to dry at room temperature.

### Real-time apoptosis and necrosis assay

For the analysis of apoptosis and necrosis, the RealTime-Glo™ Annexin V Apoptosis and Necrosis Assay (Promega) was used. 2×10^4^ HepG2 cells were seeded per well in a 96-well plate (F-bottom, white-bottomed, Greiner) in 200 µL GM and cultivated for 24 h at 37°C and 5% CO_2_ for attachment. After 24 h of incubation, GM was removed, and cells were washed with PBS. Cells were supplied with 100 µL of either phenol red-free GM or SM before addition of 100 µL of the detection reagents according to manufacturer’s instructions. Luminescence and fluorescence were measured 0, 2, 4, 6, 8, 24, and 26 h after treatment. Measurements were conducted at 37°C using orbital averaging as a scan mode with a scan diameter of 4 mm. Luminescence was measured at an emission wavelength of 470-480 nm with a gain of 3600-and 5-mm focal height. Fluorescence was measured at an excitation range of 485 ± 10 nm and collected in an emission range of 525 ± 20 with a gain of 1000 and a 5 mm focal height (CLARIOstar Plus, BMG Labtech).

### Flow cytometry ANNEXIN V/7-AAD analysi

1×106 HepG2 cells were seeded in 24-well plates (Thermo Fisher Scientific) and cultivated in GM for 24 h in a humidified atmosphere with 5% CO2 at 37°C. Subsequently, the medium was changed to either fresh GM or SM, and cells were treated with sorafenib for 24 h or 48h. Cells were then labeled according to the manual of the FITC ANNEXIN V Apoptosis Detection Kit with 7-AAD (BioLegend) with a modification where the amount of 7-AAD was doubled.

### LDH assay

Cells were seeded and treated as described for the viability assay. LDH-cytotoxicity assay kit (Takara) was used according to the manual provided, with a modification where the supernatant was diluted 10 times to stay in the measurable range of the photometer (SPARK 10M TECAN).

### Western blot

Cultured cells were scraped and collected in radioimmunoprecipitation assay buffer RIPA buffer including PhosStop and protease inhibitor cocktail (Roche). Xenograft and HCC mouse samples were homogenized in RIPA buffer using metal beads and the LT tissuelyzer (Qiagen). All samples were sonicated (ultrasound probe, 3310 s at 10%output) and centrifuged. The clear supernatant was used to measure protein concentration with a bicinchoninic acid assay (Thermo Fisher Scientific). An amount of 30 μg protein of each sample was used for western blotting. The following antibodies were used: pERK1/2 (4370), ERK1/2 (4695), pAKT (4060), AKT (4691), p4E-BP1 (2855), 4E-BP1 (9452), phospho-p70 S6 kinase (9206), p70 S6 kinase (9202), GAPDH (2118S), p21 (2947S), Bak (12105), Bax (5023), all from Cell Signaling Technologies; p53 (DO-1, sc-126, Santa Cruz Biotechnology), Vinculin (PA5-29688, Invitrogen), LC3 (NB100-2220, NOVUS), ACTB (ab6276, Abcam), and Puma (sc-374223, Santa Cruz Biotechnology). The densitometric quantification of signal intensities was performed with Image Studio Lite (Li-Cor Bioscience).

### Generating a stable p53 knock-out cell line using the CRISPR/Cas9 system

CRISPR knock-out plasmids were provided from Santa Cruz Biotechnology (sc-416469) and the experiments were performed according to the manual. 1×10^6^ HepG2 cells were seeded in a 6-well plate in antibiotic-free GM 72 h before transfection. At a confluence of 60% cells were transfected with 1,5 µg of the control or KO plasmids by using 10 µl UltraCruz transfection reagent (sc-395739). The selection of positive clones was performed by growing the cells in GM containing puromycin (sc-108071; 2µg/ml) for 10 days. Cells were analyzed and RFP+ clones were isolated via FACS (FACS Aria IIu (BD Biosciences)). Then single-cell cloning was performed to isolate RFP+ populations derived from a single cell. Different clones were transfected again (same protocol as the first transfection) with a plasmid expressing a CRE recombinase (sc-418923) and afterwards analyzed again via FACS. RFP-cells were isolated and analyzed via western blot and qPCR to validate the complete knock-out of p53.

### Overexpression of p53 by electroporation

Electroporation was performed using the Neon® Transfection System 100 μL kit (Thermo Fisher Scientific) following manufacturer’s instructions. 2×106 cells (either HepG2-WT or HepG2-KO cells) were used for each electroporation. As vectors, 2 μg of pcDNA3 flag p53 or 2 μg of HisMax as vehicle control were used. The pulse conditions voltage/width/pulses were set to 1400/30/1. The transfected cells were seeded in the appropriate cell number in 96-well plates and cultivated for 24 h in GM under standard conditions.

### RNA isolation and qPCR

RNA isolation was done using PeqGOLD Total RNA Kit according to the manufacturer’s instructions. RNA concentration was measured with NanoDrop^®^ (ND-1000, Spectrophotometer, peqLab). Isolated RNA was reverse transcribed to cDNA according to the Thermo Scientific RevertAid RT Kit using a random hexamer primer mix (1 µL). The samples (total volume of 12 µL), consisting of the template RNA (either 1 µg, 500 ng or 200 ng) diluted in nuclease-free water and the primer(s) were incubated at 65°C for 5 minutes in the DNA Engine Dyad^®^Peltier Thermal Cycler (BioRad). Afterward, the following reagents were added: 5x Reaction Buffer (4 µl), RiboLock RNase Inhibitor (20 U/µL, 1 µl), 10 mM dNTP Mix (2 µl), and RevertAid RT (200 U/µL, 1µl). The samples were then incubated in the Thermal Cycler for 5 min at 25°C, followed by 60 min at 42°C and 5 min at 70°C. The cDNA samples were diluted to 1 ng/µL and stored at - 20°C. qPCR was done in 96 or 384 well plates. Therefore, either 2.5 µL of cDNA (1 ng/µL), 5 µL of SybrGreen (BioRad) and 2.5 µL of primer-mix (800 nM, forward and reverse) per reaction in 96-well plate, or 1.5 µL of cDNA, 3 µL of SybrGreen and 2 µL of primer-mix per reaction in 384-well plate were used. Primer sequences are given in Supplementary Table 1. qPCR was performed with the CFX96™ or CFX384™ Real-Time System (C1000 Thermal Cycler, BioRad) according to the program: 1 cycle (10 min) at 95°C; 40 cycles: 15 sec at 95°C, 1 min at 60°C, 1 min at 72°C; 1 cycle: 30 sec at 95°C, 30 sec at 60°C, 30 sec at 95°C.

### Mitochondrial stress test and glycolysis stress test

To obtain confluent monolayers, 2×10^5^ cells/well were plated in five technical replicates on collagen-coated XF96 polystyrene cell culture microplates (Seahorse, Agilent) in GM, and left to adhere o/n. Cells were then incubated in GM or SM with DMSO or sorafenib as indicated in Results, and these reagents were added in all subsequent media during the measurement. For OCR measurements, cells that were kept in GM were washed and incubated in XF Base Medium (Agilent) containing 1 mM sodium pyruvate, 2 mM glutamine, and 25 mM D-glucose, and cells that were kept in SM were washed and incubated in HBSS (without Hepes) containing 5.5 mM glucose. For ECAR measurements, cells were incubated in glucose-free DMEM (D5030, Millipore Sigma) supplemented with 2 mM glutamine. An XF96 extracellular flux analyzer (Seahorse, Agilent, CA, USA) was used. After calibration of the analyzer, sequential compound injections, including oligomycin A (4 µM), FCCP (0.2 µM), antimycin A (2.5 µM), were applied to test mitochondrial respiration. Sequential compound injections, including glucose (10 mM), oligomycin A (1 µM), and 2-DG (50 mM), were applied to test glycolytic activity. OCR (pmol O_2_/min) and ECAR (mpH/min) values were normalized to protein content.

### Mitochondrial membrane potential measurements

5×10^5^ cells were seeded in GM per collagen-coated round glass in a 6-well plate and left o/n to attach. The next day, medium was changed to GM with vehicle (DMSO) or sorafenib, and cells were incubated for 2 or 6 h. Untreated cells in GM served as a control. The mitochondrial membrane potential was assessed using tetramethylrhodamine (TMRM, T668, Invitrogen) staining according to the manufacturer’s protocol. Cells were incubated with 20 nM TMRM at 37 °C for 20 min in the dark and then washed with PBS. The ∆ψ_mito_ was obtained by application of 1 µM FCCP. Imaging was performed on an inverted fluorescence microscope based on an IX73 Olympus stage (IX73 system) with a 40x Objective and a Retiga R1 CCD camera (TELEDYNE QIMAGING). Cells were excited at 550 nm and emission was captured at 600 nm.

### Mitotracker green staining

1×10^6^ cells were seeded in GM per collagen-coated round glass in a 6-well plate and left o/n to attach. The next day, medium was changed to GM with vehicle (DMSO) or sorafenib and cells were incubated for 2 or 6 h. Untreated cells were used as a control. Cells were incubated with mitotracker green (0,5 µM) (M7514, Invitrogen) in GM for 3 hours at 37°C. The mitochondrial structure was observed under an array confocal laser scanning microscope (ACLSM), based on a Zeiss Observer Z.1 inverted microscope, equipped with a YokogawaCSU-X1 Nipkow spinning disk system, a piezoelectric z-axis motorized stage (CRWG3-200; NipponThompson Co., Ltd., Tokyo, Japan), and a CoolSNAP HQ2 CCD Camera (Photometrics Tucson), using a 100× objective. The cells were excited at a wavelength of 488 nm, emission was captured at 516 nm.

### Glucose uptake measurements

1.5×10^6^ cells were seeded in GM per collagen-coated round glass in a 6-well plate and left o/n to attach. The next day, medium as changed to GM or SM with vehicle

(DMSO) or sorafenib and cells were incubated for 2 or 6 h. Untreated cells were used as a control. For Glucose uptake analysis 2-NBDG glucose-uptake marker (Thermofisher Scientific) was used. Cells were glucose depleted for 10 minutes prior measurement in loading buffer containing in mM, 2 CaCl2, 135 NaCl, 5 KCl, 1 MgCl2, 1 HEPES, 2.6 NaHCO3, 0.44 KH2PO4, 0.34 Na2HPO4, 0.1% vitamins, 0.2% essential amino acids, and 1% penicillin/streptomycin (Gibco), all at pH. After 10 minutes 2-NBDG was applied to the cells for the staining in a 0.1 mM concentration and the cells were incubated at 37°C for 30 minutes. Afterwards the cells were washed three times with a loading buffer and imaged in the same buffer. The imaging experiments were performed at an array confocal laser scanning microscope (ACLSM), based on a Zeiss Observer Z.1 inverted microscope, equipped with a YokogawaCSU-X1 Nipkow spinning disk system, a piezoelectric z-axis motorized stage (CRWG3-200; NipponThompson Co., Ltd.), and a CoolSNAP HQ2 CCD Camera (Photometrics Tucson), using a 40× objective. The glucose-uptake marker 2-NBDG was excited with a wavelength of 488 nm and the emission was captured at 516 nm. Data acquisition and control were done using the VisiView Premier Acquisition software (2.0.8, Visitron Systems). For imaging analysis, MetaMorph (Molecular Devices) was used. For the calculation of the average intensity of the cytosolic fluorescence signal, first, the ACLSM images were background-subtracted on MetaMorph using a background region of interest (ROI). MetaMorph software was applied by setting a region around the cell and performing an average intensity calculation.

### Immunohistochemistry

Immunohistochemical staining of formalin-fixed, paraffin-embedded xenografts was performed after antigen retrieval (120°C, 7 min at pH9) and peroxidase blocking (Dako) using the UltraVision LP detection system (Thermo Fisher Scientific) according to the manual with 1 ng/ml Ki-67 antibody (M728; Dako). For color reaction, AEC (3-amino-9-ethyl carbazole) chromogen (Thermo Fisher Scientific) was used. Counterstaining with hematoxylin was done on all slides. Quantitative analysis was performed by Halo® image analysis platform (Indica Labs) using Random forest tissue classifier.

### Proteomics

Mouse-derived xenograft HCC tumors were homogenized in RIPA buffer using metal beads and the LT TissueLyser (Qiagen). Samples were sonicated (Branson Sonifier 3310s ultrasound probe, 10% output), centrifuged at 14000rpm for 5min and quantified with BCA (Thermo Fisher Scientific). Cell culture (HepG2WT, HepG2KO) samples were scraped from the cell culture wells in RIPA buffer and directly sonicated, centrifuged, and quantified. 250µg of each sample was precipitated with Methanol/Chloroform protocol [84]. The protein pellet was reconstituted in 100mM Tris pH8.5, 2% sodium deoxycholate (SDC) and reduced/alkylated with 5mM TCEP/30mM chloroacetamide at 56°C for 10min. The proteins were subjected to proteolysis with 1:100 Lys-C and 1:50 trypsin overnight at 37°C. Digestion was stopped by adding 1% trifluoroacetic acid (TFA) to a final concentration of 0.5%. Precipitated SDC was removed by centrifugation at 14000rpm for 10min and supernatant containing digested peptides was desalted on Oasis HLB plate (Waters). Peptides were dried and dissolved in 2% formic acid before liquid chromatography-tandem mass spectrometry analysis. 3000 ng of the mixture tryptic peptides were analyzed using an Ultimate3000 high-performance liquid chromatography system (Thermo Fisher Scientific) coupled on-line to a Q Exactive HF-x mass spectrometer (Thermo Fisher Scientific). Buffer A consisted of water acidified with 0.1% Formic Acid, while Buffer B was 80% Acetonitrile (ACN) and 20% water with 0.1% FA. The peptides were first trapped for 1 minute at 30 µL/min with 100% buffer A on a trap (0.3 x 5mm with PepMap C18, 5μm - 100Å Thermo Fisher Scientific); after the trapping peptides were separated by a 50-cm analytical column packed with C18 beads (Poroshell 120 EC-C18, 2.7 μm, Agilent Technologies). The gradient was 9-40%B in 155 minutes at 400 nL/min. Buffer B was then raised to 55% in 10 minutes and increased to 99% for the cleaning step. Peptides were ionized using a spray voltage of 1.9 kV and a capillary heated at 275 °C. The mass spectrometer was set to acquire full-scan MS spectra (350–1400 m/z) for a maximum injection time of 120 ms at a mass resolution of 120,000 and an automated gain control (AGC) target value of 3×10^6^. Up to 25 of the most intense precursor ions were selected for tandem mass spectrometry (MS/MS). HCD fragmentation was performed in the HCD cell, with the readout in the Orbitrap mass analyzer at a resolution of 15,000 (isolation window of 1.4 Th) and an AGC target value of 1×10^5^ with a maximum injection time of 25 ms and a normalized collision energy of 27%.

### Proteomics data analysis

All raw files were analyzed by MaxQuant v1.6.17 software using the integrated Andromeda Search engine and searched against the Human UniProt Reference Proteome (October 2020 release with 75,088 protein sequences) alone for cells, while for xenograft we added the Mouse UniProt Reference Proteome (October 2020 release with 55,489 protein sequences) [85]. MaxQuant was used with the standard parameters (the “Label-Free Quantification” and “Match between runs” were selected with automatic values) with only the addition of Deamidation (N) as variable modification. Data analysis was then carried out with Perseus v1.6.14: proteins reported in the file “proteinGroups.txt” were filtered for reverse, potential contaminants and identified by site. For the quantitation we used the LFQ calculated by MaxQuant and we kept only proteins found in at least 2 biological replicates in each group; in xenografts we eliminated proteins which were only identified in the Mouse Fasta. At this point missing values were imputed by Perseus with the automatic settings (width: 0.3, down shift: 1.8, mode: separately for each column) leading to 5,064 proteins left for statistical analysis with ANOVA testing (Benjamini-Hochberg FDR 0.05), z-score (mean per row) and hierarchical clustering. The mass spectrometry proteomics data have been deposited to the ProteomeXchange Consortium via the PRIDE [86] partner repository with the dataset identifier PXD022723.

### In vivo experiments

All experiments were performed in accordance with the European Directive 2010/63/EU and approved by the Austrian Federal Ministry of Education, Science and Research. Mice were housed under standard 12 h light/12 h dark cycles. Sorafenib was first dissolved in DMSO (120 mg/ml) and then diluted in PEG400:PBS (1:1) in a final working concentration of 3 mg/ml. Mice received 30 mg/kg sorafenib by oral gavage.

### Xenograft assays

2×10^6^ HepG2 WT or KO cells in 100 µL PBS:Matrigel (1:1) were injected into both hind flanks of 6 week old male NMRI-*Foxn1*^*nu*^ mice. Once the tumors were measurable (>100 mm^3^, 14-21 days after injection), the mice were randomly assigned to 4 groups (n=6-10 per group) receiving either vehicle control (DMSO in PEG 400:PBS (1:1)) or sorafenib via oral gavage in groups that were kept on ad libitum diet (Altromin 1320 fortified) or on an intermittent fasting regimen, meaning that mice were withheld food for 24 hours two times a week (with ad libitum access to water and hydrogel), with two- or three-days ad libitum refeeding in-between. Sorafenib or vehicle control was applied at the beginning of the refeeding phase and at a concentration of 30 mg/kg. Xenografts were measured with a caliper once a week and tumor volume was calculated using a formula: V = (4/3) × π × (L/2) × (L/2) × (W/2), L-length (shorter dimension), W-width (longer dimension). At the end of the fourth IF cycle, mice were refed for 3-4 days and then sacrificed. Some mice were sacrificed after an additional 24 h fast.

### Double transgenic mice

Tp53^*flox/flox*^ C57BL/6J mice were crossed with CreER^*T2+/-*^ C57BL/6J mice to generate Tp53^*flox/flox*^ CreER^*T2+/-*^ mice and the respective Tp53^*flox/flox*^ CreER^*T2-/-*^ controls [59].

### Pilot experiments

Pilot experiments were performed to establish the most effective intermittent fasting protocol and safe sorafenib concentrations. In the first pilot experiment C57BL/6J mice (n=6 per group) were fasted for 24 h with 48h of refeeding in-between, while in the second pilot experiment C57BL/6J mice (n=7 per group) were fasted every other day for one week. Food intake and body weight were measured daily. For NMR measurements, blood was collected from the submandibular vein immediately before animals were sacrificed, after the last fasting day. Between 100 and 300 µl of blood was mixed with 10 µL EDTA and kept at room temperature for a maximum of 30 min. Then, samples were centrifuged at 4°C, at 3500 rpm for 10 min. Plasma was transferred to clean tubes and stored at -80°C until measurement.

A third pilot experiment was performed to assess the liver toxicity of sorafenib. C57BL/6J mice (n=6 per group) received vehicle, 10, 30 or 50 mg/kg sorafenib by oral gavage three times per week for 4 consecutive weeks. Mice were weighed two times per week (when gavaged). At the end of the experiment, mice were sacrificed and liver pieces were formalin-fixed and paraffin-embedded for further analyses.

### NMR metabolite analysis

To extract polar metabolites, 140 µl of ice-cold methanol were added to 70 µl of plasma, the tube mixed by shaking, and stored at - 20°C for 1h. Tubes were centrifuged at 13000 rpm for 30 min (4°C), the supernatant transferred to a fresh tube, and the samples lyophilized. For NMR experiments, samples were re-dissolved in 500 µl of NMR buffer [0.08 M Na2HPO4, 5 mM TSP (3-(trimethylsilyl) propionic acid-2,2,3,3-d4 sodium salt), 0.04 (w/v) % NaN3 in D2O, pH adjusted to 7.4 with 8 M HCl and 5 M NaOH]. Plasma metabolic analysis was conducted at 310 K using a Bruker Advance Neo 600 MHz NMR spectrometer equipped with a TXI probe head. The Carr–Purcell–Meiboom–Gill (CPMG) pulse sequence was used to acquire 1H 1D NMR spectra with pre-saturation for water suppression (cpmgpr1d, 512 scans, 73728 points in F1, 12019.230 Hz spectral width, 1024 transients, recycle delay 4 s). NMR spectral data were processed as previously described [87]. Shortly, data were processed in Bruker Topspin version 4.0.2 using one-dimensional exponential window multiplication of the FID, Fourier transformation and phase correction. NMR data were then imported to Matlab2014b, TSP was used as an internal standard for chemical shift referencing (set to 0 ppm), regions around the water, TSP and methanol signals were excluded, NMR spectra were aligned, and a probabilistic quotient normalization was performed. Quantification of metabolites was carried out by signal integration of normalized spectra.

### DEN experiment

N-Nitrosodiethylamine (DEN, N0756 Sigma-Aldrich), was diluted in 0.9% sterile NaCl to a working concentration of 2.5 mg/ml. Male CreER^T2+/-^ and CreER^T2-/-^ Tp53^flox/flox^ C57BL/6J mice received 25 mg/kg DEN i.p. at the age of two weeks. After 41-43 weeks, after developing HCC, the mice received 100 mg/kg tamoxifen by oral gavage for five consecutive days, followed by a one-week wash-out phase. Tamoxifen (Molekula) was prepared fresh as a 30 mg/ml working solution in 90% peanut oil and 10% ethanol. Then, four cycles of treatment were conducted: mice were either fed ad libitum (FED) or fasted two times per week for 24 h in the intermittently fasted (IF) group. Both groups received 30 mg/kg sorafenib by oral gavage two times per week, after the 24 h fasting in IF group, and on the same day in FED group. After four cycles, and additional 2-3 days of ad libitum feeding, the mice were sacrificed.

### Ultrasound

Briefly, mice were anesthetized using isoflurane (2 L/min O_2_, 4% isoflurane in the anesthesia chamber; 2 L/min O_2_, 2% isoflurane in the anesthesia mask during imaging). Anesthetized animals were shaved and fixed lying down on the working bench. Ultrasound imaging was performed using Vevo 3100 (FUJIFILM VisualSonics).

### Liver biochemistry

Liver panel strips (Menarini SPOTCHEM II, Liver-1 33925) were used for the quantitative determination of LDH, GPT, GOT, albumin, total protein and total bilirubin in mouse plasma (by Spotchem EZ SP-4430 Arkray Inc.). Briefly, blood was collected from the submandibular vein immediately before animals were sacrificed. Between 100 and 300 µl of blood was mixed with 10 µL EDTA and left at room temperature for a maximum of 30 min. Then, samples were centrifuged at 4°C, at 3500 rpm for 10 min. Plasma was transferred to clean tubes and stored at -80°C until measurement. For the panel measurements, 70 µl of plasma was loaded.

### Statistics

If not stated otherwise, all experiments were performed at least three times independently. Statistical analyses were performed using GraphPad Prism 8 (GraphPad Software). Statistically significant differences were determined as described in figure legends. If not noted otherwise, data represent mean values ± SEM with the following grades of statistical significance: *p <0.05, **p <0.01, ***p <0.001. Statistical proteomic analyses (ANOVA, z-score, hierarchical clustering) were performed using Perseus v1.6.14.0 [88].

## AUTHOR CONTRIBUTIONS

J.K. designed and performed the experiments, analyzed the data, and wrote the manuscript. I.R., K.S., N.B., N.K., E.M., M.A., H.M., and C.N. performed experiments and analyzed data. M.R.D., J.R., and S.S. performed and analyzed experiments on cell metabolism. M.G. and R.Z. performed and analyzed proteomics experiments. M.P., B.R., A.J.A.D, A.R., T.M., A.J.R.H., M.H., and R.M. contributed materials and provided expertise and feedback. A.P. designed, coordinated, and supervised the project, analyzed the data and wrote the manuscript.

## ACKNOWLEDGEMENTS

We thank Sandra Blass, Helmut Bischof, Stefanie Wallner, Robert Arnes-Benito, Ivan Vidakovic, and Ruth Prassl for technical and methodological help. We are grateful to Mitch Lazar, Michael Schupp, Roberto Flores, Ariane Pessentheiner, Ellen Heitzer, and Sasa Frank for critical evaluation of the manuscript. We also thank Prof. Pierre Chambon, Prof. Richard Moriggl, and Doris Kaltenecker for making Alb-CreERT2 mice available to us and Prof. Tim Schulz for providing p53-floxed mice. We are grateful to Prof. Berthold Huppertz for providing support at the Division of Cell Biology, Histology, and Embryology. J.K. was funded and supported by the Austrian Science Fund (FWF, grant P29328). I.R. was funded by the PhD faculty MolMed at the Medical University of Graz. I.R., M.G., K.S., N.B., N.K., H.M., and C.N. were supported by the Austrian Science Fund (FWF, grants P29328 and I3165). A.P. was supported by the Austrian Science Fund (FWF, grants P29328 and I3165) and by a MEFO grant from the Medical University of Graz. This work has also been supported by EPIC-XS, project number 823839, funded by the Horizon 2020 program of the European Union. T.M. was supported by Austrian Science Foundation Grants P28854, I3792, and DK-MCD W1226; Austrian Research Promotion Agency (FFG) Grants 864690 and 870454; the Integrative Metabolism Research Center Graz; Austrian Infrastructure Program 2016/2017, the Styrian Government (Zukunftsfonds), and BioTechMed-Graz (Flagship project). S.S., M.G. N.B., and N.K. were trained within the frame of the PhD program Molecular Medicine, Medical University of Graz.

