## Supplemental Figures and Table for "Fasting reverses drug-resistance in hepatocellular carcinoma through p53-dependent metabolic synergism"

### Figure S1

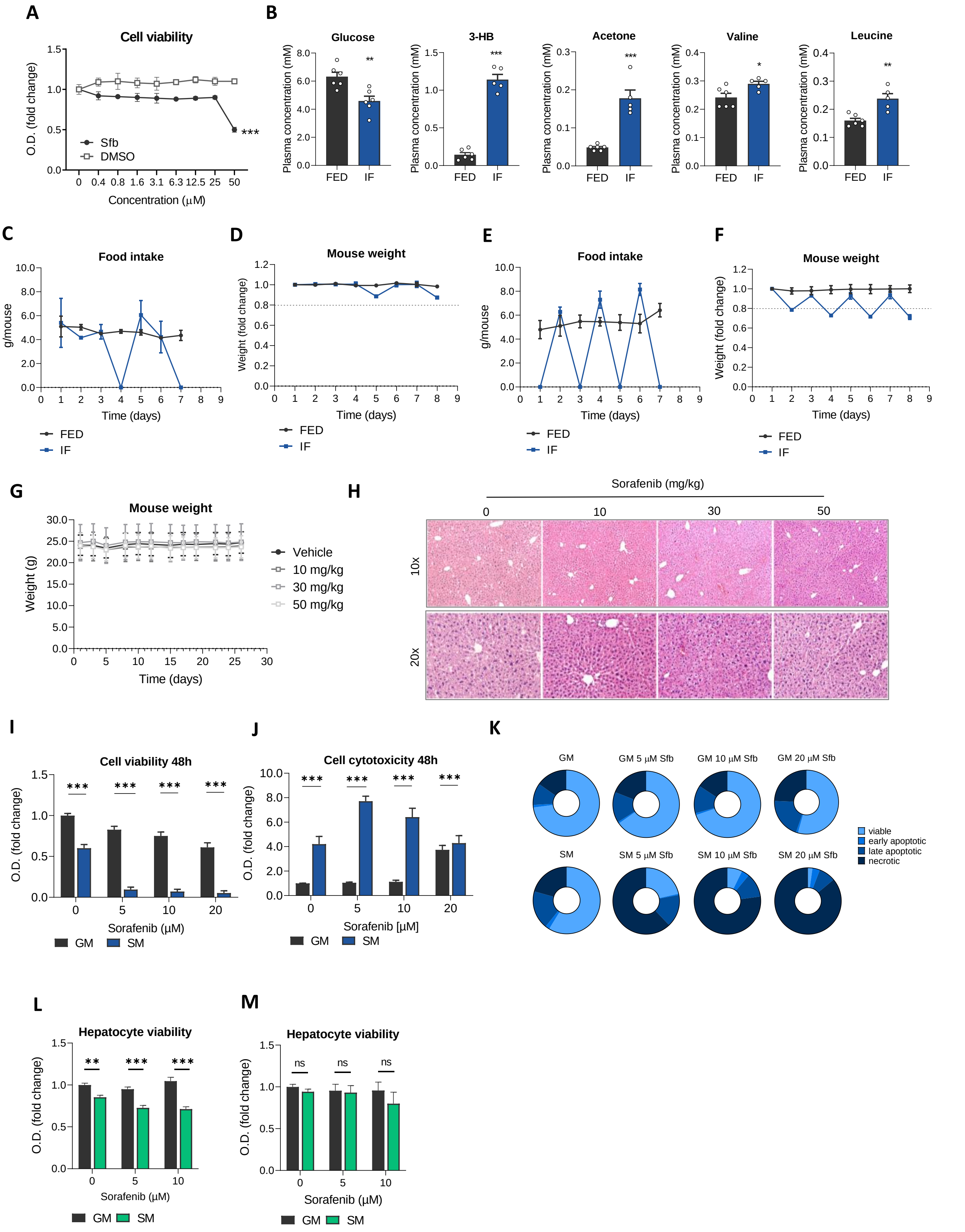
A
Cell viability
O.D. (fold change)
Concentration (μM)
Sfb
DMSO
***
B
Glucose
3-HB
Acetone
Valine
Leucine
Plasma concentration (mM)
FED
IF
C
Food intake
g/mouse
Time (days)
D
Mouse weight
Weight (fold change)
E
Food intake
F
Mouse weight
G
Mouse weight
Weight (g)
Vehicle
10 mg/kg
30 mg/kg
50 mg/kg
H
Sorafenib (mg/kg)
0
10
30
50
10x
20x
I
Cell viability 48h
Sorafenib (μM)
GM
SM
J
Cell cytotoxicity 48h
Sorafenib [μM]
K
GM
GM 5 μM Sfb
GM 10 μM Sfb
GM 20 μM Sfb
SM
SM 5 μM Sfb
SM 10 μM Sfb
SM 20 μM Sfb
viable
early apoptotic
late apoptotic
necrotic
L
Hepatocyte viability
M
Hepatocyte viability
ns

**Figure S1**.
(A) HepG2 cell viability after 24h incubation with sorafenib (Sfb) or corresponding volumes of vehicle (DMSO). The viability at each timepoint was normalized to untreated cells at zero time. Two-way ANOVA, Dunnett's multiple comparisons test, all values compared to control, untreated cells.
(B) NMR analysis of metabolites in mouse plasma after 24h fasting. Two-sided student's t-test was performed.
Pilot experiments were performed to establish the most effective intermittent fasting and sorafenib protocol. In the first pilot experiment C57BL/6J mice were fasted for 24h with 48h of refeeding in-between:
(C) Daily food intake ± SD.
(D) Mean normalized weights ± SD of ad libitum fed (FED) and intermittently fasted (IF) mice. Mice lost less than 20% of their starting body weight during 24h fasting and regained their weight during 48h refeeding.
In the second pilot experiment C57BL/6J mice were fasted every other day:
(E) Daily food intake ± SD.
(F) Normalized mean weights ± SD of FED and IF mice. Mice lost more than 20% of their starting body weight by the end of this protocol.
Third pilot experiment was performed to assess the liver toxicity of sorafenib. C57BL/6J mice (n=6 per group) received sorafenib by oral gavage three times per week in designated concentrations for 4 consecutive weeks:
(G) Mean mouse weight ± SD.
(H) Hematoxylin and eosin staining of liver sections after 4 weeks of treatment.
(I) Viability of HepG2 cells after 48 h of incubation in GM or SM with indicated concentrations of Sfb. Values are normalized to the viability of non-treated cells grown in GM.
(J) Cytotoxicity measured by LDH assay in cells grown in GM vs cells grown in SM with indicated Sfb concentrations after 48h.
(K) Proportions of viable (7AAD and Annexin V negative), early apoptotic (Annexin V positive), late apoptotic (Annexin V and 7AAD positive) and dead (7AAD positive) cells analyzed by flow cytometry after 24-hour incubation in GM or SM with indicated sorafenib concentrations.
(L) Mouse primary hepatocytes were cultured for 18h in GM, followed by 6h in SM with or without sorafenib. Then a viability assay was performed.
(M) Mouse primary hepatocytes were cultured in GM or SM with or without sorafenib for 6h. Then a viability assay was performed.
If not noted otherwise, mean values ± SEM are shown and two-way ANOVA, Tukey's multiple comparisons test was performed. *** $p<0.001$, ** $p<0.01$, * $p<0.05$, ns-not significant ($p>0.05$); O.D. optical density; nd-not detected, O.D. optical density.

### Figure S2

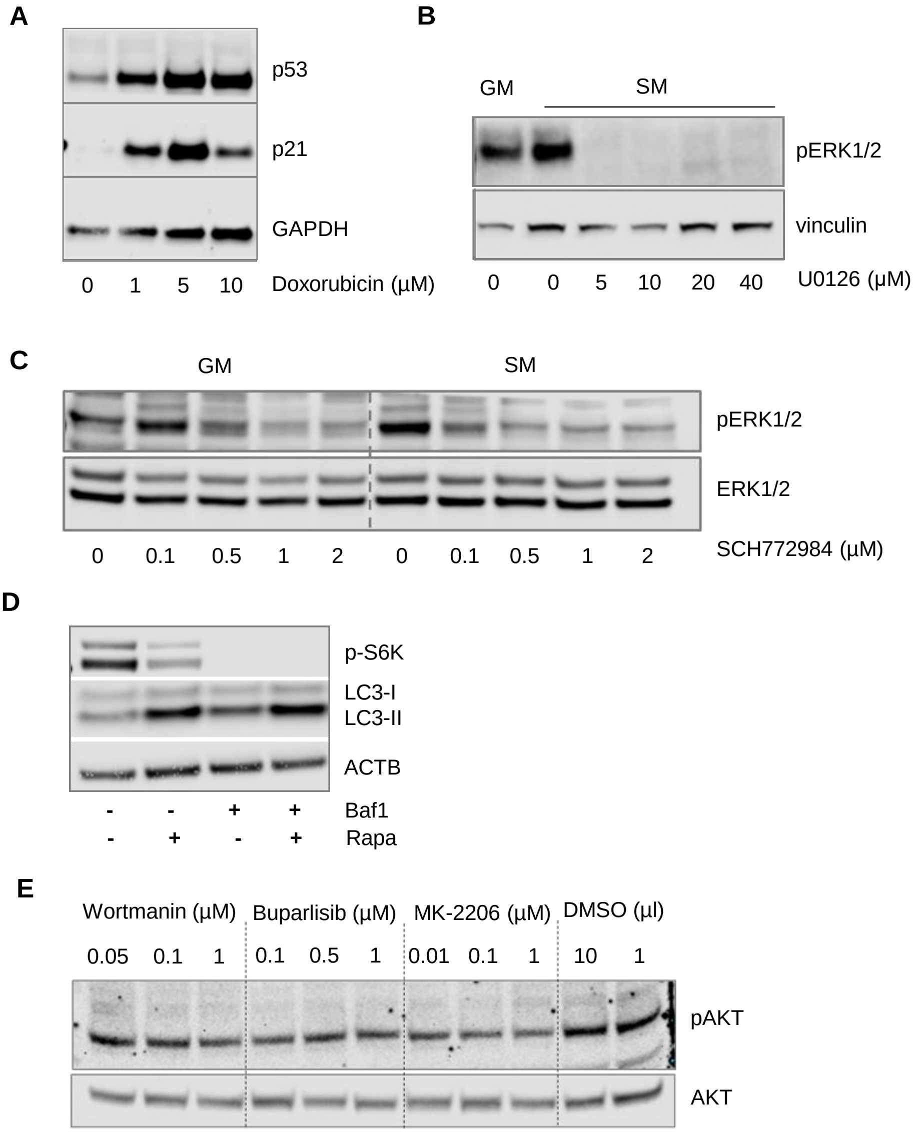


**Figure S2.**

**(A)** Stabilization of p53 and p21 in HepG2 cells after 24h of treatment with doxorubicin determined by western blot. GAPDH served as a loading control.

**(B)** Inhibition of ERK1/2 phosphorylation by MEK inhibitor, U0126, demonstrated by western blot. HepG2 cels were grown in growth medium (GM) or starvation medium (SM) with or without UO126 for 24h.

**(C)** HepG2 cells were grown in GM or in SM with or without ERK1/2 inhibitor, SCH772984, for 24h. pERK1/2 inhibition was demonstrated by western blot.

**(D)** HepG2 cells were treated with 10 nM bafilomycin A1 and/or 100 nM rapamycin for 2 hours. Western blotting shows increased lipidated LC3 (LC3-II) upon bafilomycin A1 treatment (indicating blocking of autophagy) and complete inhibition of mTOR signalling by rapamycin.

**(E)** Inhibition of AKT phosphorylation via designated inhibitors determined by western blot. Cell lysates were prepared after 24h treatment with each of the inhibitors in GM. Control cells were treated with corresponding volumes of the vehicle (DMSO).

### Figure S3

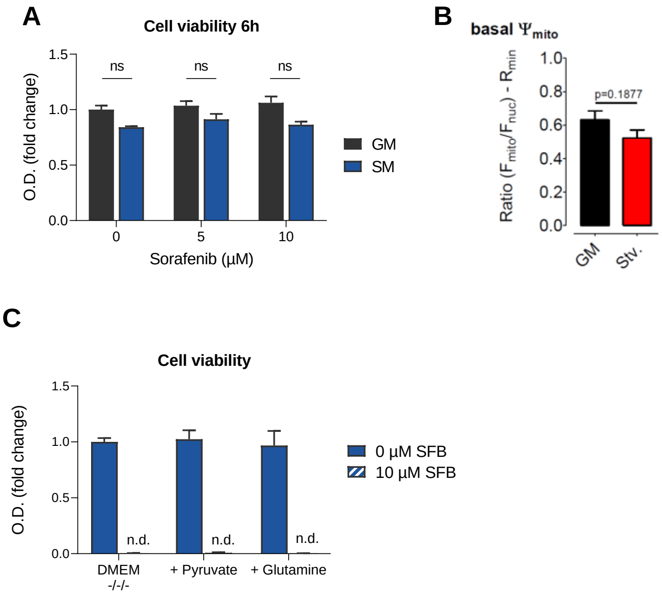


**Figure S3.**
**(A)** HepG2 cells were kept in growth medium (GM) or starvation medium (SM) with indicated concentrations of sorafenib (Sfb) for 6h. Then the viability assay was performed.
**(B)** Mitochondrial membrane potential measured using TMRM in HepG2 cells after 6h incubation in GM and SM.
**(C)** HepG2 cells were kept in DMEM without glucose, glutamine and pyruvate ($DMEM^{-/-/-}$) or with added 4mM glutamine or 100 mM sodium pyruvate and with and w/o sorafenib. After 24h viability assay was performed.
If not noted otherwise, mean values ± SEM are shown and two-way ANOVA, Tukey's multiple comparisons test was performed. *** p<0.001, ns-not significant (p>0.05); O.D. optical density.

### Figure S4

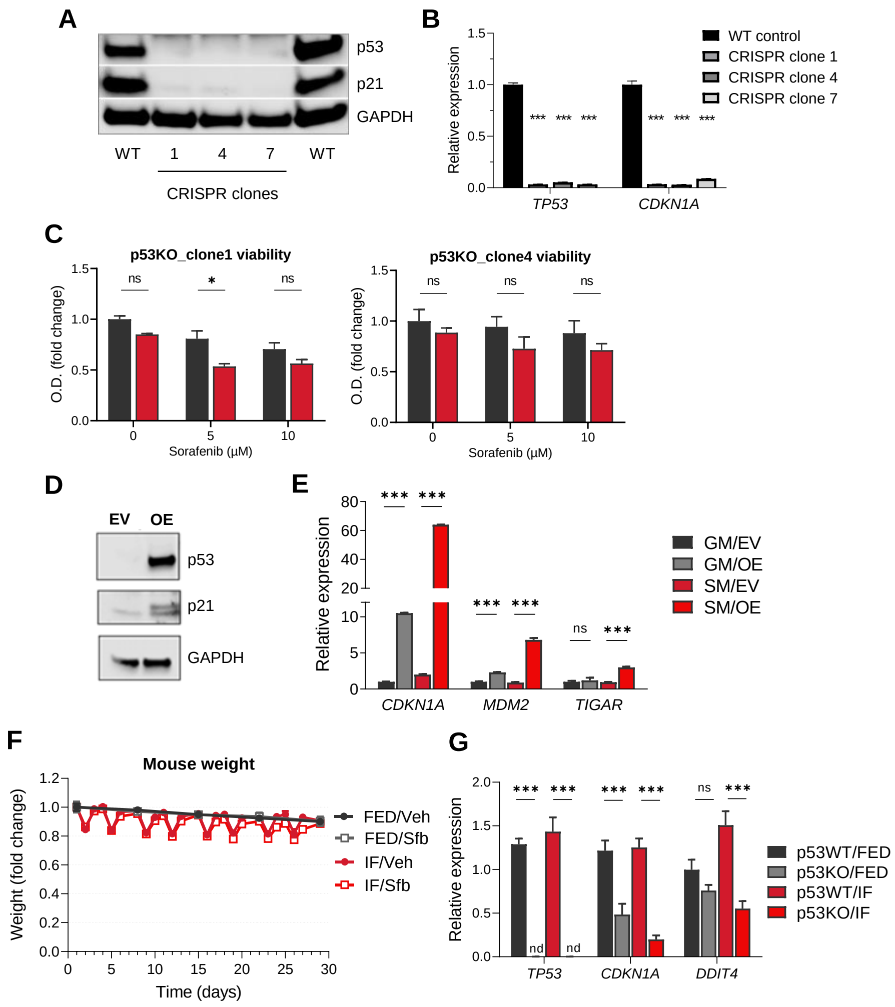


**Figure S4.**
**(A)** Western blot analysis and
**(B)** qPCR analysis was performed to asses the p53 knock-out in HepG2 cells.
**(C)** Two additional p53KO HepG2 clones were kept in growth medium (GM) or starvation medium (SM) with 0, 5, 10 and 20 µM sorafenib. Viability assay was performed after 24h of incubation and normalized to the viability of non-treated cells grown in GM. Viability of cells grown in GM vs cells grown in SM was analyzed.
**(D)** Western blot was performed with cell lysates from p53KO cells transfected with empty vector (EV) or p53 overexpression vector (OE) to determine p53 and p21 levels. GAPDH served as loading control.
**(E)** qPCR analysis of p53 target gene expression, 48h after p53 knock-out.
**(F)** Animal whole body weight during the treatment, normalized to starting weight of each group. Mean values ± SD are shown; n=8-10 in each group.
**(G)** qPCR analysis was performed to assess *TP53* and target gene expression in HepG2 xenografts at the end of treatment.
If not noted otherwise, mean values ± SEM are shown and two-way ANOVA, Tukey's multiple comparisons test was performed. *** p<0.001, ** p<0.01, * p<0.05, ns-not significant (p>0.05); O.D. optical density.

### Figure S5

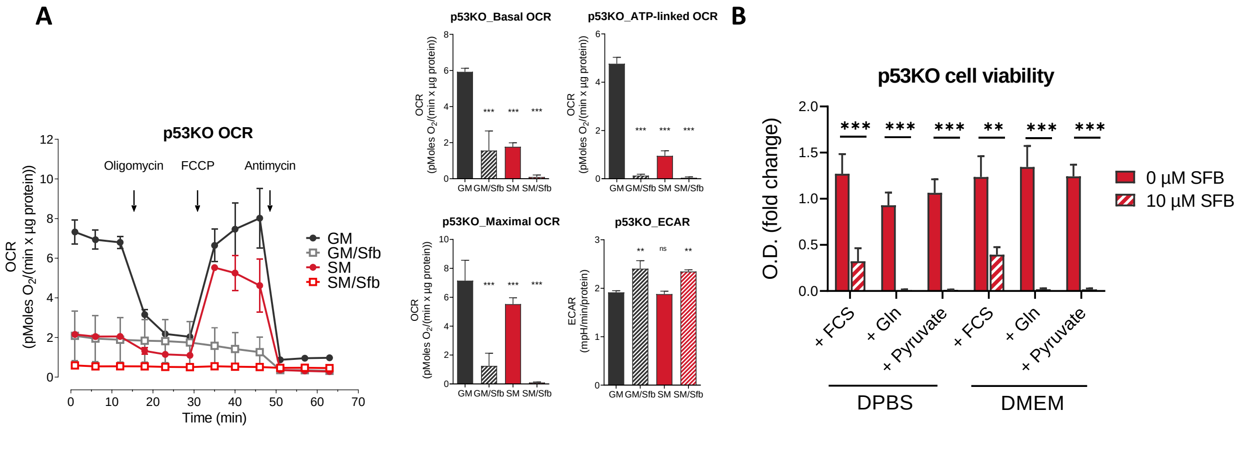


**Figure S5.**

**(A)** Seahorse analysis of mitochondrial respiration in p53KO HepG2 cells. Cells were incubated for 6 hours in growth medium (GM) or starvation medium (SM) with 0 and 10 µM sorafenib. Cells were then incubated 1h prior experiment in XF assay medium supplemented with 5 mM glucose and 2 mM glutamine and consecutively injected with oligomycin (4 µM), FCCP (0.2 µM), and antimycin (2.5 µM). Continuous and basal oxygen consumption rate (OCR) values normalized to protein content in cells are shown.

**(B)** Confluent p53KO HepG2 cells were grown in DPBS or DMEM-/-/- with the addition of 10% FCS, 4mM glutamine or 100 mM sodium pyruvate. After 24h viability assay was performed.

If not noted otherwise, mean values ± SEM are shown and two-way ANOVA, Tukey's multiple comparisons test was performed. *** $p<0.001$, ** $p<0.01$, * $p<0.05$, ns-not significant ($p>0.05$); O.D. optical density.

### Figure S6

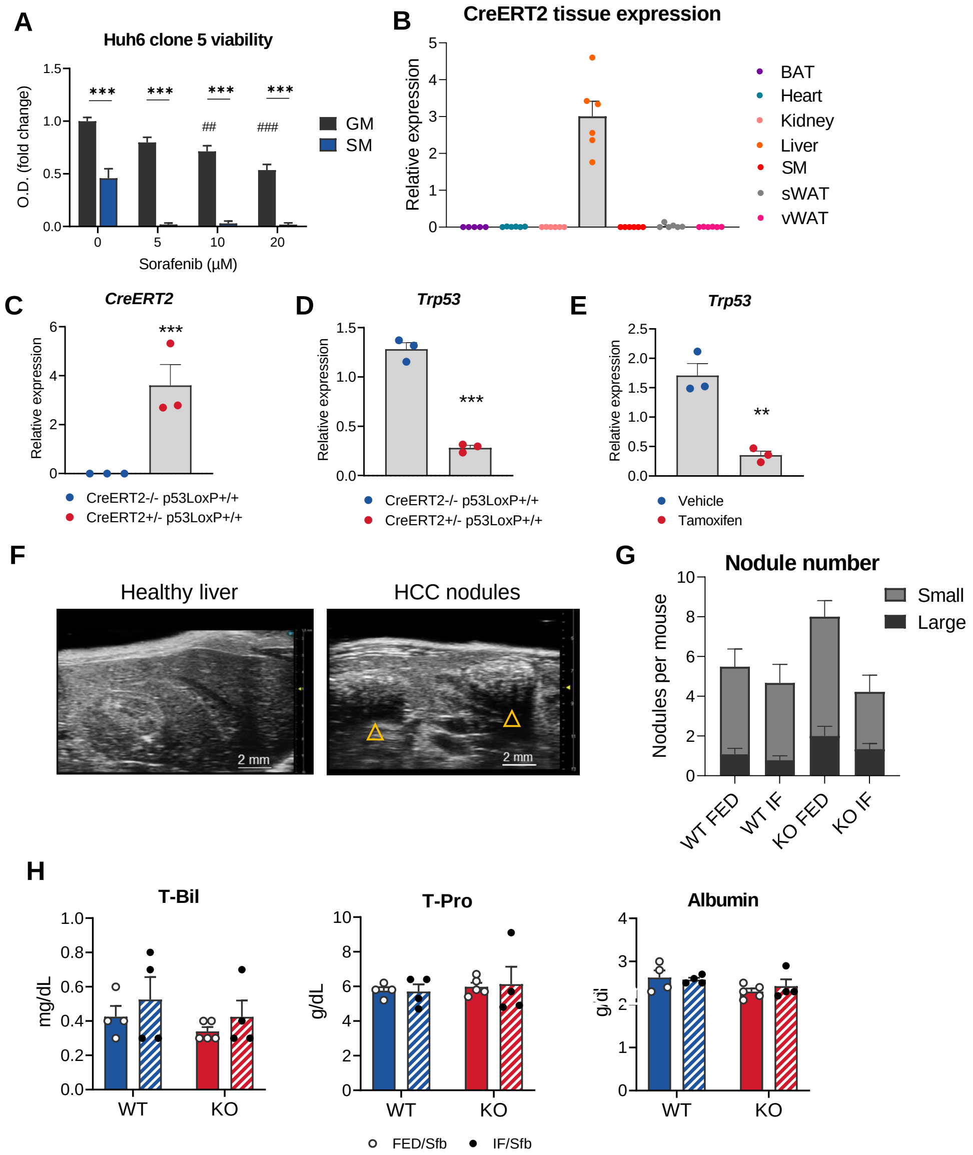


**Figure S6**. Pilot experiments were performed to assess the p53 knock-out in double transgenic mice. qPCR analyses were performed to analyze:

(A) Huh6 clone5 cell line was kept in growth medium (GM) or starvation medium (SM) with 0, 5, 10, and 20µM sorafenib (Sfb). Viability assay was performed after 24 hours of incubation. Comparison between GM and SM groups (*), and vs control, GM group (#) is shown

(B) the expression of CreERT2 in CreER$^{T2+/-}$ Tp53$^{flox/flox}$ mice treated with 100 mg/kg tamoxifen for five consecutive days, followed by 14-day wash-out phase.

(C) the expression of CreERT2 and

(D) Trp53 in the liver of CreER$^{T2+/-}$ Tp53$^{flox/flox}$ and CreER$^{T2-/-}$ Tp53$^{flox/flox}$ mice treated with 100 mg/kg tamoxifen for five consecutive days, followed by 14-day wash-out phase. Unpaired t-test was performed.

(E) the expression of Trp53 in CreER$^{T2+/-}$ Tp53$^{flox/flox}$ mice treated with vehicle or 100 mg/kg tamoxifen for five consecutive days, followed by 14-day wash-out phase. Unpaired t-test was performed.

(F) Ultrasound images of HCC nodules in mice 33 weeks after DEN i.p.

(G) Small (<3mm) and large (>3mm) HCC nodule number at the end of experiment.

(H) Concentrations of T-Bil, T-Pro and Albumin in the plasma on the sacrifice day in mice from designated groups.

If not noted otherwise, mean values ± SEM are shown and Two-way ANOVA, Tukey's multiple comparisons test was performed. *** p<0.001, ** p<0.01, * p<0.05, ns-not significant (p>0.05).

### Table S1

| | |
|---|---|
| **Human primers 5'-3'** | |
| SLC2A1_F | TCTGGCATCAACGCTGTCTTC |
| SLC2A1_R | CGATACCGGAGCCAATGGT |
| SLC2A3_F | GCTGGGCATCGTTGTTGGA |
| SLC2A3_R | GCACTTTGTAGGATAGCAGGAAG |
| TP53_F | CAGCACATGACGGAGGTTGT |
| TP53_R | TCATCCAAATACTCCACACGC |
| CDKN1A_F | GGCAGACCAGCATGACAGATT |
| CDKN1A_R | GCGGATTAGGGCTTCCTCTT |
| MDM2_F | GAATCATCGGACTCAGGTACATC |
| MDM2_R | TCTGTCTCACTAATTGCTCTCCT |
| TIGAR_F | ACTCAAGACTTCGGGAAAGGA |
| TIGAR_R | CACGCATTTTCACCTGGTCC |
| DDIT4_F | TGAGGATGAACACTTGTGTGC |
| DDIT4_R | CCAACTGGCTAGGCATCAGC |
| BAK1_F | ATGGTCACCTTACCTCTGCAA |
| BAK_R | TCATAGCGTCGGTTGATGTCG |
| BAX_F | TTTTGCTTCAGGGTTTCATC |
| BAX_R | CAGTTGAAGTTGCCGTCAGA |
| PUMA_F | CTGTGAATCCTGTGCTCTGC |
| PUMA_R | AATGAATGCCAGTGGTCACA |
| GAPDH_F | ACCCACTCCTCCACCTTTGA |
| GAPDH_R | CTGTTGCTGTAGCCAAATTCGT |
| PPIA_F | GGCAAATGCTGGACCCAACACA |
| PPIA_R | TGCTGGTCTTGCCATTCCTGGA |
| B2M_F | CCACTGAAAAAGATGAGTATGCCT |
| B2M_R | CCAATCCAAATGCGGCATCTTCA |
| **Mouse primers 5' - 3'** | |
| Slc2a1_F | CATCCTTATTGCCCAGGTGTT |
| Slc2a1_R | GAAGACGACACTGAGCAGCAG |
| Slc2a2_F | CCCTGGGTACTCTTCACCAA |
| Slc2a2_R | GCCAAGTAGGATGTGCCAAT |
| Slc2a3_F | TCATCTTCGCTGCCTTCCT |
| Slc2a3_R | CAGCACTCAGAAGCAGTCCTGGT |
| Slc2a5_F | GCTGCAGCCAAATTGCCCAAT |
| Slc2a5_R | CGGGGCCAGCTCCCCTAAGT |
| Bak1_F | TGCCTACGAACTCTTCACCAA |
| Bak1_R | TGGTAGACGTACAGGGCCAG |
| Bax_F | TGAAGACAGGGGCCTTTTTG |
| Bax_R | AATTCGCCGGAGACACTCG |
| Puma_F | CGACCTCAACGCGCAGTA |
| Puma_R | AGTCCCATGAAGAGATTGTACATGAC |
| Trp53_F | ACATGACGGAGGTCGTGAG |
| Trp53_R | AATTTCCTTCCACCCGGATA |
| CreERT2_F | CGGTCTGGCAGTAAAAACTAT |
| CreERT2_R | CAGGGTGTTATAAGCAATCCC |
| Ppia_F | CAGACGCCACTGTCGCTTT |
| Ppia_R | TGTCTTTGGAACTTTGTCTGCAA |
| B2m_F | ATGGCTCGCTCGGTGACCCT |
| B2m_R | TTCTCCGGTGGGTGGCGTGA |
